# Atypical MEG inter-subject correlation during listening to continuous natural speech in dyslexia

**DOI:** 10.1101/677674

**Authors:** A. Thiede, E. Glerean, T. Kujala, L. Parkkonen

## Abstract

Listening to speech elicits brain activity time-locked to the speech sounds. This so-called neural entrainment to speech was found to be atypical in dyslexia, a reading impairment associated with neural speech processing deficits. We hypothesized that the brain responses of dyslexic vs. normal readers to real-life speech would be different, and thus the strength of inter-subject correlation (ISC) would differ from that of typical readers and be reflected in reading-related measures.

We recorded magnetoencephalograms (MEG) of 23 dyslexic and 21 typically-reading adults during listening to ∼10 min of natural Finnish speech consisting of excerpts from radio news, a podcast, a self-recorded audiobook chapter and small talk. The amplitude envelopes of band-pass-filtered MEG source signals were correlated between subjects in a cortically-constrained source space in six frequency bands. The resulting ISCs of dyslexic and typical readers were compared with a permutation-based *t*-test. Neuropsychological measures of phonological processing, technical reading, and working memory were correlated with the ISCs utilizing the Mantel test.

During listening to speech, ISCs were reduced in dyslexic compared to typical readers in delta (0.5–4 Hz), alpha (8–12 Hz), low gamma (25–45 Hz) and high gamma (55–90 Hz) frequency bands. In the beta (12–25 Hz) band, dyslexics had mainly enhanced ISC to speech compared to controls. Furthermore, we found that ISCs across both groups were associated with phonological processing, technical reading, and working memory.

The atypical ISC to natural speech in dyslexics supports the temporal sampling deficit theory of dyslexia. It also suggests over-synchronization to phoneme-rate information in speech, which could indicate more effort-demanding sampling of phonemes from speech in dyslexia. These irregularities in parsing speech are likely some of the complex neural factors contributing to dyslexia. The associations between neural coupling and reading-related skills further support this notion.

**Research Highlights:**

- MEG inter-subject correlation (ISC) of dyslexics was atypical while listening to speech.
- Depending on the frequency band, dyslexics had stronger or weaker ISC than controls.
- Reading-related measures correlated with the strength of ISC.

## 1 Introduction

Language processing and comprehension are essential for human communication and interaction. Neural speech processing deficiencies are typical for individuals with developmental dyslexia, a learning disorder characterized by reading and writing difficulties affecting up to 17% of the population (Elliott and Grigorenko, 2014). The speech processing deficit in dyslexia has been investigated widely (for reviews, see e.g. Ramus et al., 2003; Schulte-Körne and Bruder, 2010), however, mostly by utilizing unnatural, repetitive stimuli that barely resemble real-life speech. It has been argued that to truly understand the mechanisms of language processing in real-life situations, naturalistic stimuli should be used (Hasson et al., 2018). The core question of this study is whether the neural dynamics of processing natural speech are atypical in dyslexia.

This question has previously been illuminated from different angles. For example, acoustic and rhythmic properties of the speech stimulus *per se* are reflected in oscillatory brain activity, which has been suggested to enhance speech perception and comprehension (Doelling et al., 2014; Luo and Poeppel, 2007; Obleser and Weisz, 2012; Peelle and Davis, 2012), differently so in dyslexics than typical readers (De Vos et al., 2017a; Power et al., 2016). The natural brain rhythms (i.e., oscillations) thereby seem to interplay with the speech stimulus that is being processed (for a review, see Meyer, 2018). One interesting aspect, however, has not gained much attention in the field of speech processing in dyslexia: Brain synchronization. When incoming information, such as speech, is processed in a similar manner across individuals, their neural activity is likely synchronized as well, which leads to a common understanding and goal-directed behaviour (Hasson et al., 2012). The extent of synchronization can be estimated with inter-subject correlation (ISC), a model-free analysis approach that has been proven viable to extract shared brain activations across participants during natural stimulation due to the time-varying dynamics of the stimulus (Hasson et al., 2004). ISC has been extensively applied during naturalistic paradigms in fMRI, e.g. movie viewing (Hasson et al., 2004; Jääskeläinen et al., 2008; Kauppi et al., 2010; Nummenmaa et al., 2012), music listening (Abrams et al., 2013; Alluri et al., 2013), and speech processing (Wilson et al., 2008; Stephens et al., 2010; Lerner et al., 2011; Silbert et al., 2014; Finn et al., 2018). However, its application to MEG has been a lot more scarce. The only MEG ISC studies to date have looked at movie viewing with various ISC methodologies (Suppanen, 2014; Lankinen et al., 2014; Chang et al., 2015) and music listening (Thiede, 2014). The scarcity of MEG ISC studies could arise from the non-trivial methodology (e.g. complexity of the MEG signal, ill-posed source estimation problem), lack of ISC implementations for MEG as well as the substantial computational power required to do ISC analysis with MEG data. However, compared to fMRI, MEG can reveal new, complementary information that enables addressing slightly different questions. Whereas fMRI measures brain activity indirectly through the sluggish hemodynamic response and can only track fluctuations < 1 Hz, MEG directly measures electric activity of neuronal populations with millisecond resolution. FMRI is also more affected by blood-oxygenating physiological processes in the body, e.g. pulsation and breathing.

The richness of the MEG signal allows extracting several measures (e.g. phase coupling, envelope correlation, cross-frequency coupling) across different frequency bands during rest or task. We focus here on one aspect; the envelope correlation in a set of frequency bands while the subject is listening to speech. ISC reflects functioning of cortical areas that respond to the time-varying stimulus dynamics, which in speech are manifold: For example, acoustic, phonological, syntactic, and semantic features likely activate lower- and higher-level brain functions related to processing and comprehension of speech. In functional magnetic resonance imaging (fMRI) studies, ISCs were found in healthy adult participants listening to natural speech in bilateral temporal areas, frontal areas, parietal areas including premotor cortex, and midline areas including precuneus (Wilson et al., 2008; Stephens et al., 2010; Lerner et al., 2011; Silbert et al., 2014; Finn et al., 2018). The first objective of the current study was to confirm and extend our knowledge of the brain areas that couple between healthy adult participants during listening to natural speech using magnetoencephalography (MEG).

Certain brain dynamics have been repeatedly shown to be abnormal in dyslexia, specifically during speech processing. For example, temporal sampling deficits have been proposed to play a role in dyslexia, especially in the delta and theta band which reflect syllable encoding (Goswami, 2011; Hämäläinen et al., 2012; Molinaro et al., 2016). Moreover, Giraud and Poeppel (2012) have proposed that speech parsing at rates comparable to low-gamma frequencies is altered in dyslexia. Indeed, brain measures during processing of speech correlate with reading-related tests. For example, an abnormal right-rather than left-lateralized auditory steady-state response in dyslexics was associated with behavioural tests of phonology, and further, a phonemic oversampling, i.e. faster than normal oscillatory rate, has been associated with memory deficits in dyslexia (Lehongre et al., 2011). The second objective of the present study was to investigate whether brain activity of dyslexics during listening to speech is atypically synchronized compared to typical readers. We hypothesized that especially lower frequency bands (Goswami, 2011; Hämäläinen et al., 2012; Molinaro et al., 2016) show weaker ISCs between dyslexic than typical readers, whereas higher frequency bands could show enhanced ISCs between dyslexic compared to typical readers (Lehongre et al., 2011). Thirdly, we examined the association between ISC and neurophysiological measures across both groups. We hypothesized that the strength of ISC is associated with reading-related test performance.

These hypotheses were assessed by comparing the ISCs of MEG amplitude envelopes during listening to natural speech in dyslexic and typical readers. The MEG amplitude envelopes were extracted in the cortically-constrained source space of each individual in six frequency bands of interest (delta, theta, alpha, beta, low gamma, high gamma). Then, pairwise correlations were computed and averaged to obtain group correlations that were compared between groups. We found significant differences in ISC to speech between the groups, and could further show that the strength of ISC was associated with reading-related skills. These results reveal atypical processing of natural speech in dyslexia and show that these brain dynamics are reflected in reading-related skills.

## 2 Methods

This study has been preregistered at ClinicalTrials.gov (<u>NCT02622360</u>) as part of a research project on speech- and short-term memory functions in dyslexia.

### 2.1 Participants

Forty-nine Finnish-speaking right-handed adult participants aged 18–45 years and without a history of neurological diseases volunteered in the study, 26 with confirmed dyslexia and 23 typical readers. Participants were recruited from an organization for learning impairments (HERO Ry, Helsinki, Finland) as well as from university and adult education email lists, from a related project website, and by an advertisement in social media. To be included in the dyslexic group, participants had to have 1) a diagnosis from a psychologist, special education teacher, or similar, 2) evident reading-related problems in childhood indicated by the adult reading history questionnaire (ARHQ; Lefly and Pennington, 2000) and confirmed in an interview, and 3) below-norm performance (less than one standard deviation from the age-matched average) in at least two reading subtests in either speed or accuracy (see Section 2.2). To be included in the control group, 1) participants or their relatives had to have no language-related disorders, 2) the ARHQ indicated no reading-related problems in childhood, and 3) participants had to perform within norm in at least two reading subtests. Exclusion criteria for the study were attention deficits (ADD) as tested by the Adult ADHD Self-Report Scale ASRS-v1.1 questionnaire (Kessler et al., 2005), other language impairments, such as developmental language disorder (formerly specific language impairment), other neurological or psychiatric diseases, medication severely affecting the central nervous system, a special education track in school indicative of wider cognitive impairments, non-compensated hearing or sight deficits, and a performance intelligence quotient (IQ) below 80. Data of four participants were excluded as anatomical MRIs could not be obtained due to metal in the body or pregnancy (three dyslexics, one control), and data from one participant had to be excluded due to technical reasons during the MEG measurement which resulted in missing trigger markers (control). The final sample consisted of 44 participants, of which 23 were in the dyslexic and 21 in the control group. Background information are summarized in Table 1; statistics were performed with SPSS version 24.0 (IBM Corp, 2016, Armonk, NY, USA). Participants gave their written consent after they had been informed about the study. All procedures were carried out according to the Declaration of Helsinki, and the Coordinating Ethics Committee (Hospital District of Helsinki and Uusimaa) approved the study protocol.

**Table 1.** Descriptive statistics about background information regarding both groups (dyslexic, control) and statistics for group differences. For scalar variables (age, education and musical education), means (M, bold) and standard deviations (STD) are reported and independent-sample t-tests are used for group difference statistics. For the categorical variable (gender), the count for each category (male/female, m/f) is reported and the Χ^2^-test is used for group difference statistics.

| VARIABLE | DYSLEXIC GROUP |  |  | CONTROL GROUP |  |  | STATISTICS |  |  |
| --- | --- | --- | --- | --- | --- | --- | --- | --- | --- |
|  | *N* | *M* | *STD* | *N* | *M* | *STD* | *t/X<sup>2</sup>* | *df* | *p* |
| AGE [YEARS] | 23 | **31.6** | 8.7 | 21 | **30.0** | 6.0 | 0.71 | 42 | .482 |
| GENDER [COUNT] | 23 | 11/12 | (m/f) | 21 | 10/11 | (m/f) | 1.89E-04 | 1 | .989 |
| EDUCATION [YEARS] | 23 | **15.7** | 5.2 | 20 | **17.0** | 2.6 | -0.95 | 41 | .347 |
| MUSICAL EDUCATION [YEARS] | 23 | **3.0** | 7.8 | 21 | **3.1** | 4.8 | -0.04 | 42 | .972 |

### 2.2 Neuropsychological tests

Neuropsychological tests were conducted by Master students of psychology under the supervision of a licensed clinical psychologist in a session of ca. 2 h at the Cognitive Brain Research Unit, University of Helsinki. Domains of phonological processing, reading, IQ, and memory functions were assessed. Phonological processing was evaluated with the ‘Pig Latin’ test (Nevala et al., 2006), non-word span length (Laasonen et al., 2002), digit span length (Wechsler, 2008), and rapid alternating stimulus naming (Wolf, 1986). Reading skills were evaluated by word and pseudoword list reading (technical reading) and text reading (reading comprehension; Nevala et al., 2006). The verbal IQ was assessed with similarities and vocabulary subtests, and performance IQ with block design and matrix reasoning subtests (Wechsler, 2005). Memory function was evaluated with the subtests on letter-number series and visual series (Wechsler, 2008). A summary of the neuropsychological test outcomes is presented in Table 2; statistics were performed with SPSS, effect sizes were calculated with Psychometrica Freeware (Lenhard, 2017, Dettelbach, Germany, https://www.psychometrica.de/effect_size.html#cohenb, https://www.psychometrica.de/effect_size.html#nonparametric), and bootstrapped confidence intervals were calculated with the measures-of-effect-size toolbox (Hentschke and Stüttgen, 2011, https://github.com/hhentschke/measures-of-effect-size-toolbox). Composite scores were formed for phonological processing and technical reading by converting the raw scores to z-scores and averaging them, and for working memory the composite was formed according to WMS-III (Wechsler, 2008).

**Table 2.**
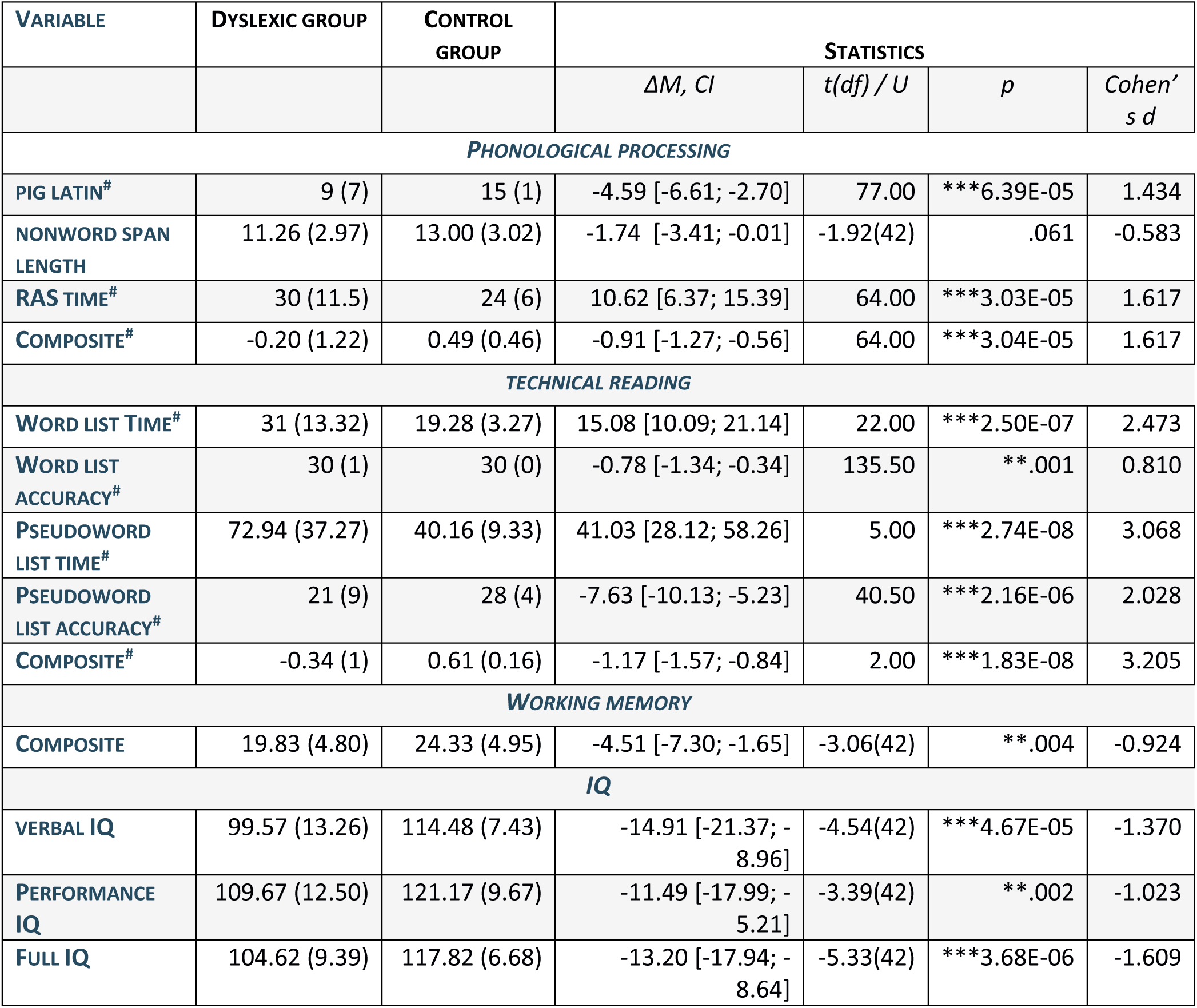
Descriptive statistics on neuropsychological test performances for both groups (dyslexic, control). Reported are means, standard deviations (in brackets), mean differences (ΔM) with bootstrapped confidence intervals (CI), t-values with degrees of freedom (df, in brackets) and p-values of group comparisons from independent-sample t-tests, and Cohen’s d effect sizes for normally distributed scores in both groups. For non-normally distributed scores in one or both groups(^#^), median, interquartile range (in brackets), mean differences (ΔM) with bootstrapped confidence intervals (CI), U-values and p-values of group comparisons from Mann-Whitney U-tests, and Cohen’s d effect sizes are reported. FDR-corrected significance levels are marked with asterisks (*p < 0.046, **p < 0.01, ***p < 0.001). Composite scores were formed for phonological processing and technical reading by converting the raw scores to z scores and averaging them, and for working memory the composite was formed according to WMS-III (Wechsler, 2008).

| VARIABLE | DYSLEXIC GROUP | CONTROL GROUP | STATISTICS |  |  |  |
| --- | --- | --- | --- | --- | --- | --- |
| | | | $\Delta M$, CI | t(df) / U | p | Cohen's d |
| **PHONOLOGICAL PROCESSING** |  |  |  |  |  |  |
| PIG LATIN<sup>#</sup> | 9 (7) | 15 (1) | -4.59 [-6.61; -2.70] | 77.00 | ***6.39E-05 | 1.434 |
| NONWORD SPAN LENGTH | 11.26 (2.97) | 13.00 (3.02) | -1.74 [-3.41; -0.01] | -1.92(42) | .061 | -0.583 |
| RAS TIME<sup>#</sup> | 30 (11.5) | 24 (6) | 10.62 [6.37; 15.39] | 64.00 | ***3.03E-05 | 1.617 |
| COMPOSITE<sup>#</sup> | -0.20 (1.22) | 0.49 (0.46) | -0.91 [-1.27; -0.56] | 64.00 | ***3.04E-05 | 1.617 |
| **TECHNICAL READING** |  |  |  |  |  |  |
| WORD LIST TIME<sup>#</sup> | 31 (13.32) | 19.28 (3.27) | 15.08 [10.09; 21.14] | 22.00 | ***2.50E-07 | 2.473 |
| WORD LIST ACCURACY<sup>#</sup> | 30 (1) | 30 (0) | -0.78 [-1.34; -0.34] | 135.50 | **0.001 | 0.810 |
| PSEUDOWORD LIST TIME<sup>#</sup> | 72.94 (37.27) | 40.16 (9.33) | 41.03 [28.12; 58.26] | 5.00 | ***2.74E-08 | 3.068 |
| PSEUDOWORD LIST ACCURACY<sup>#</sup> | 21 (9) | 28 (4) | -7.63 [-10.13; -5.23] | 40.50 | ***2.16E-06 | 2.028 |
| COMPOSITE<sup>#</sup> | -0.34 (1) | 0.61 (0.16) | -1.17 [-1.57; -0.84] | 2.00 | ***1.83E-08 | 3.205 |
| **WORKING MEMORY** |  |  |  |  |  |  |
| COMPOSITE | 19.83 (4.80) | 24.33 (4.95) | -4.51 [-7.30; -1.65] | -3.06(42) | **0.004 | -0.924 |
| **IQ** |  |  |  |  |  |  |
| VERBAL IQ | 99.57 (13.26) | 114.48 (7.43) | -14.91 [-21.37; -8.96] | -4.54(42) | ***4.67E-05 | -1.370 |
| PERFORMANCE IQ | 109.67 (12.50) | 121.17 (9.67) | -11.49 [-17.99; -5.21] | -3.39(42) | **0.002 | -1.023 |
| FULL IQ | 104.62 (9.39) | 117.82 (6.68) | -13.20 [-17.94; -8.64] | -5.33(42) | ***3.68E-06 | -1.609 |

### 2.3 Stimuli and data acquisition

Natural Finnish speech of ≈10 min was used as the auditory stimulus (sampling rate 44100 Hz; original sound file, transcription and its translation to English in Supplementary Material). The stimulus consisted of several shorter excerpts that were merged into one audio file with Audacity® 2.0 software (Audacity Team, 2012, http://audacityteam.org/). All excerpts were spoken by native Finnish speakers and either extracted from online sources (Finnish national broadcast ‘Yle’ radio news and podcast) or recorded by the experimenters (reading a book and small talk, such as asking for directions and exchanging of travel experiences) in a sound-proof laboratory at the Cognitive Brain Research Unit, University of Helsinki. The excerpts were chosen to represent a wide range of voices (male and female), topics, and style (conversation, factual, lyrical). Consecutive excerpts were joined with a 1-s silent break with 0.5-s fade-out and 0.5-s fade-in. The waveform of the speech stimulus is visualized in Figure 1A.

**Figure 1.**
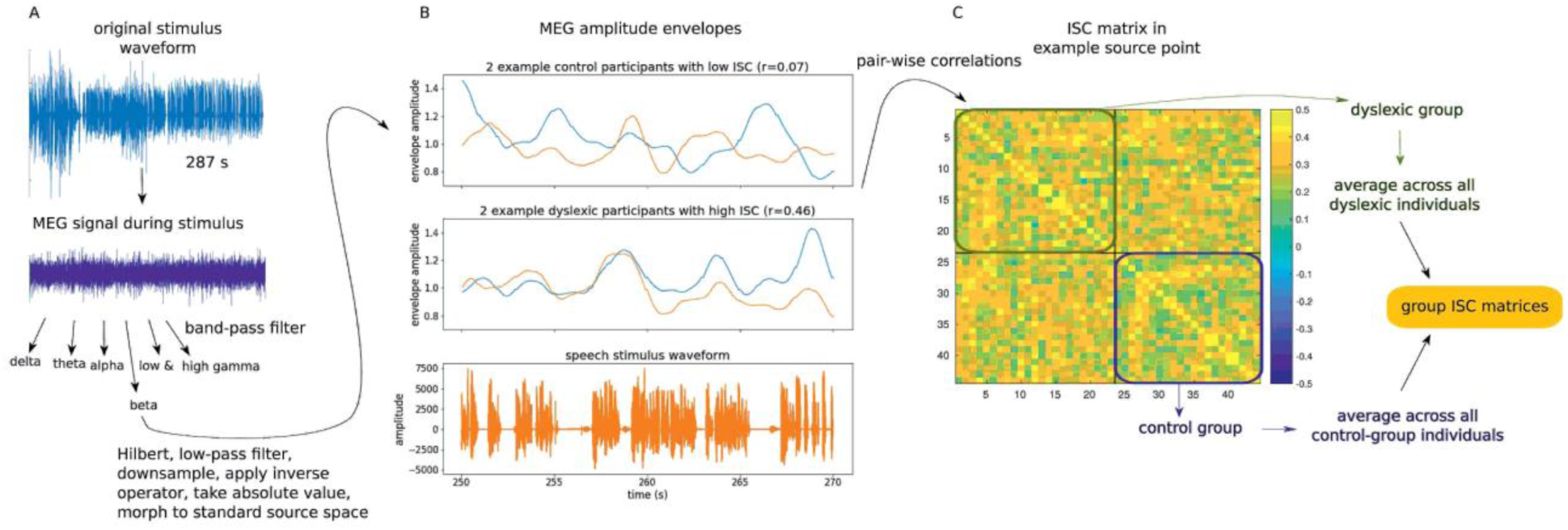
Schematic representation of the inter-subject correlation (ISC) data analysis. *A.* Acoustic waveform of the speech stimulus (part 1, duration 287 s). The MEG signal was extracted during the time of the stimulus. Here, the preprocessed MEG signal of an example channel (MEG1622) above the left temporal area is shown. The MEG signal was then filtered to six frequency bands (delta, theta, alpha, beta, low gamma, high gamma), Hilbert-transformed, low-pass filtered, downsampled, source modelled, and finally the absolute value was taken to obtain the instantaneous amplitude at every source point and in all six frequency bands. The source locations of these amplitude signals were then morphed from individual cortical source space to a standard source space. *B.* Beta-band MEG amplitude envelopes of example participants showing low ISC (top panel) and high ISC (middle panel) at a source in the middle temporal cortex. The waveform of the speech stimulus during the same excerpt of 20 s is shown for comparison (bottom panel). *C.* ISC matrix of all pairwise correlations at the same source location as in B). The upper left square (olive frame) contains ISC values for dyslexic pairs and the bottom right square (blue frame) for control pairs. Group ISC matrices were obtained at all source points by averaging across all individuals of one group.

The neural activity of the brain was recorded with an Elekta Neuromag Triux MEG system (MEGIN Oy, Helsinki, Finland) comprising 204 planar gradiometers and 102 magnetometers. The signals were filtered to 0.03–330 Hz and sampled at 1 kHz. Recordings were performed in a magnetically shielded room (Euroshield/ETS Lindgren Oy, Eura, Finland) at BioMag Laboratory in Helsinki University Hospital. Participants listened to the continuous auditory stream binaurally at a comfortable level (≈70–80 dB SPL). The stimulus was presented with Presentation Software (Neurobehavioral Systems Ltd., Berkeley, CA, USA) and conveyed from earphones to the ears via plastic tubes. Resting-state MEG data (eyes open) were recorded for each participant for ≈10 min. Other auditory and visual stimuli (written pseudowords and the corresponding auditory versions as well as scrambled visual symbols) had been presented before these recordings for ≈80 min in six recording blocks. Data from these recordings will be presented in separate publications. In all MEG recordings, participants were seated in an upright position and were instructed to relax and to listen to the continuous speech stimulus while keeping the head still.

In addition to MEG, scalp EEG and horizontal and vertical electrooculograms (EOG) were recorded with a 60-channel cap (EasyCap, Herrsching, Germany) with reference and ground electrodes located at the nose and left cheek, respectively. Five head position indicator coils (HPI), the EEG electrodes, and fiducial markers of nasion and both preauricular points were digitized with a Polhemus Isotrak 3D-digitizer (Polhemus Inc., Colchester, VT, USA) in order to establish a transformation between the MEG and MRI coordinate systems. The HPI coils were continuously energized to enable tracking and compensation of head movements throughout the MEG measurement.

Structural T1-weighted magnetic resonance images (MPRAGE sequence) were obtained with a 3T MAGNETOM Skyra whole-body MRI scanner (Siemens Healthcare, Erlangen, Germany) with a standard 32-channel head coil at AMI centre, Aalto University. Each structural MRI consisted of 176 slices with a slice thickness of 1 mm, voxel size of (1 x 1 x 1) mm^3^, and field of view of (256 x 256) mm^2^. All structural MRIs were checked by a physician who reported no incidental findings.

### 2.4 Data analysis

The code used for the analysis of this dataset is available at https://github.com/athiede13/free_speech.

#### MEG data preprocessing

The continuous MEG data were preprocessed by first visually examining all recordings and marking noisy, flat, or otherwise artifact-containing channels as bad (on average 6.2 channels in one recording). External magnetic interference was suppressed with Maxfilter software version 2.2 (MEGIN Oy, Helsinki, Finland) applying temporal signal-space separation (tSSS; Taulu and Simola, 2006) with a buffer length of 10 s and correlation limit of 0.98. The algorithm also corrected for head movements measured with the HPI coils and interpolated the channels manually marked or automatically detected as bad. Physiological artifacts, specifically those resulting from eye blinks, eye movements, and heartbeats, were removed with signal-space projection (SSP; Tesche et al., 1995; Uusitalo and Ilmoniemi, 1997) implemented in MNE-Python (Gramfort et al., 2014; 2013) software package (<u>version 0.17.dev0</u>). Channels that showed the most prominent artifacts (EOG channels for eye-movements and channel ‘MEG1541’ for heartbeats) were used to average the artifact events and create the projectors. The noise covariance was estimated with MNE-Python from ‘empty-room’ data of ≈10 min that were preprocessed similarly to the data from the participants.

#### MRI data preprocessing

Structural MRIs were preprocessed using the Freesurfer software package (versions 5.3 and 6.0, Martinos Center for Biomedical Imaging, http://freesurfer.net/; Dale et al., 1999; Fischl et al., 1999a, 1999b). The steps applied included segmentation of brain volume with the watershed algorithm (Ségonne et al., 2004), intensity normalization (Sled et al., 1998), segmentation of grey and white matter (Fischl et al., 2004, 2002), and inflation of the cortical surfaces (Fischl et al., 1999a). Manual editing of surfaces, performed by an experienced graduate student, was required in 66% of the cases to ensure a correct segmentation of the brain volume and manual addition of white-matter points in 18% to ensure a correct segmentation of the grey and white matter boundary.

#### Coregistration

Coregistration of MRI and MEG was performed with the function *mne coreg* in the MNE-Python software package. First, the digitized fiducials and head-shape points (EEG electrode positions) were manually aligned with the reconstructed head surface from the individual anatomical MRI. Then, the iterative closest point algorithm was applied to minimize the distances of the head-shape points from the head surface.

#### Source modeling

The segmented cortical surface was decimated (recursively subdivided octahedron) to yield 4098 source points per hemisphere. A single-compartment boundary-element model (BEM) was applied to compute the forward solution; source points closer than 5 mm to the BEM surface were omitted. A dSPM minimum-norm estimate (MNE) inverse operator was then computed with a loose orientation constraint of 0.2, depth weighting exponent of 0.8, and the noise covariance estimated from the ‘empty-room’ data.

#### Inter-subject correlation (ISC)

For ISC computation (for an overview, see Figure 1), custom scripts were utilized in MATLAB (release 2017a; The MathWorks, Inc., Natick, Massachusetts, USA) as well as the MNE Matlab toolbox (Gramfort et al., 2014) and MEG ISC custom functions (Suppanen, 2014; Thiede, 2014). First, in the listening-to-speech condition, the stimulus durations and temporal alignments with respect to the recordings were determined with the help of the stimulus start and end triggers from Presentation (due to technical reasons, the stimulus was in two parts; 4.77 and 5.45 min). For the determined stimulus durations, the preprocessed MEG signals were band-pass filtered (third-order Butterworth filter, applied in the forward direction only) into six frequency bands of interest (cut-off frequencies; delta: 0.5–4 Hz, theta: 4–8 Hz, alpha: 8–12 Hz, beta: 12–25 Hz, low gamma: 25–45 Hz, high gamma: 55–90 Hz). The analytical signals were computed by applying Hilbert transformation to the band-pass-filtered signal. The resulting signals were low-pass filtered (similar filter as above) at 0.3 Hz, and downsampled to 10 Hz. The previously computed inverse operator was then applied to these complex-valued signals. The absolute value of each source time series was taken, resulting in cortical amplitude envelopes per each participant and frequency band (delta, theta, alpha, beta, low gamma, high gamma). The cortical locations of the envelopes were morphed from each individual subject to the Freesurfer standard brain (*fsaverage*) with MNE-Python. The source space of this standard brain consists of 20484 points per hemisphere, causing an automatic upsampling of the source points during the morphing step. Pairwise correlations of the cortical amplitude envelopes at the corresponding source points were computed across all subject pairs within each experiment group and for each frequency band. The pairwise correlations were averaged for each group, i.e., dyslexic and control group. A duration-weighted averaging was applied for the two speech parts.

To test whether ISCs were significantly larger than zero, a spatial permutation-based one-sample *t*-test was applied to the group-average ISC matrices (MNE-Python function *spatio_temporal_cluster_1samp_test* based on Maris & Oostenveld (2007)). First, this test calculates the statistic (one-sample *T*-test) and forms initial clusters that are above the threshold using spatial neighborhood information; second, it permutes the data by randomized sign flips (subject pair labels are permuted here), finds clusters from each permutation, and returns the maximal cluster sizes; third, it returns clusters and corrected *p*-values that are computed as a percentile of the statistic within the ‘null distribution’ taken from the surrogate data generated by the permutations. The initial *p*-threshold for cluster formation was 0.05, the *t*-threshold was 1.97, and the number of permutations was 5000. The spatial connectivity was estimated from the *fsaverage* source space including all immediate neighbors. *T*-values of clusters that survived the cluster-*p*-threshold of 0.05/6 (Bonferroni-correction for the six frequency bands) were visualized.

The ISC contrast between the groups was then tested with a spatial permutation -based *t*-test with 5000 permutations (function *spatio_temporal_cluster_test* in MNE-Python). This test differs from the above one in the first step: Here, an independent-samples *T*-test was calculated, and the *t*-threshold of ±1.97 was considered for contrasts in both directions. The permutation test was performed for both negative and positive thresholds separately, and the positive and negative clusters that survived the cluster-*p*-threshold of 0.05/6 (Bonferroni-correction for the six frequency bands) were combined and visualized.

#### Correlation between ISC strengths and neuropsychological tests

We tested for correlations between the brain-to-brain coupling strength during listening to speech (ISCs) and neuropsychological test scores using the Mantel test (Mantel, 1967). The neuropsychological test scores were combined into four composite measures: phonological processing, technical reading, working memory, and IQ (see Section 2.2).

Computations were carried out with custom scripts in MATLAB and MNE Python. Regression matrices were computed as models for the Mantel test by averaging the test scores between each subject pair for all four neuropsychological composites. Surrogate maps were computed by random permutation of the subject labels for 5000 times. The Mantel test was performed as a Spearman rank correlation between the top triangle of the ISC matrix (all pairwise combinations) and the top triangle of the regression matrix reflecting the neuropsychological composite (four composites of interest: phonological processing, technical reading, working memory, IQ). The ISC matrix contained values for each subject pair (946 pairs) and source location (20484 locations), and an uncorrected *p*-value was estimated for each source with the Mantel test. An uncorrected *r*-threshold was computed for each frequency band.

Cluster correction was performed by finding clusters for each surrogate map (5000) that exceeded the uncorrected *r*-threshold using the spatial connectivity information. For each model, the maximal cluster size was returned; the 5000 values represented the null distribution of cluster sizes. We then adopted the maximum statistics approach to control for all comparisons across all frequency bands (Winkler et al., 2016). From the surrogate maps obtained with permutations, the maximum of all maximal cluster sizes across frequencies and neuropsychological composites (24 computations) was computed as a cutoff for the real Mantel data. Clusters were formed in the same way for the real Mantel data as for the surrogate maps, and only clusters larger than the cutoff size were visualized on the *fsaverage* brain provided by Freesurfer.

To showcase the distribution of correlation between each neuropsychological composite and ISC for control and dyslexic pairs, the mean ISC in the largest cluster was plotted against the corresponding composite scores for each frequency band.

## 3 Results

### 3.1 Interbrain correlation during listening to speech

ISCs were significantly larger than zero in all frequency bands and in both groups and exhibited different correlation strengths across frequency bands (Figure 2, Table 3). Two large clusters encompassing the two complete hemispheres (with 10242 source locations in each) were found, because of the spatial spreading of the L2 MNE and the large number of sample pairs in the correlation computation.

**Figure 2.**
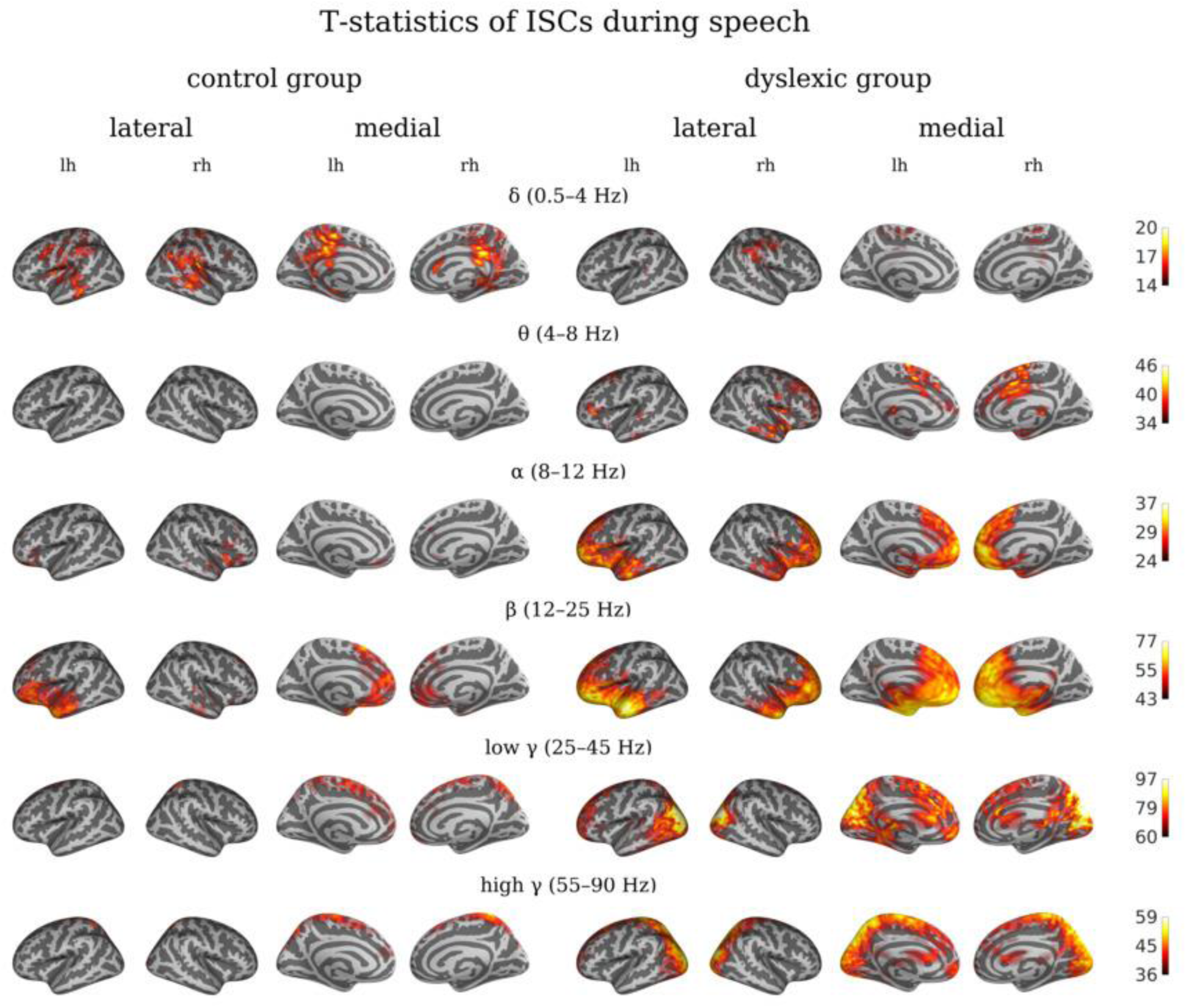
T-statistics of permutation-based one-sample t-tests for inter-subject correlations (ISCs) during listening to speech in control (left four views) and dyslexic (right four views) group. ISCs are depicted in six MEG frequency bands (delta, theta, alpha, beta, low gamma, high gamma) in lateral (first two views of each group) and medial (last two views of each group) views (lh – left hemisphere, rh – right hemisphere).

**Table 3.** . Peak MNI coordinates in significant frequency bands, cluster sizes, t/r-statistic (maximum/minimum of the largest cluster), and corresponding automated anatomical labeling (AAL) brain area (Brodmann area, BA, in brackets) for 1) ISC clusters during listening to speech for both groups, 2) ISC brain areas with group differences (con - control group, dys - dyslexic group), and 3) brain areas with significant regression between ISCs during listening to speech and reading-related measures.

| frequency band | cluster size | MNI coordinates (x, y, z) |  |  | t/r | AAL brain area (BA) |
| --- | --- | --- | --- | --- | --- | --- |
| 1) ISC > 0 |  |  |  |  |  |  |
| CONTROL GROUP |  |  |  |  |  |  |
| delta | 10242 | 4 | -31 | 30 | 20.57 | Cingulum_Mid_R (23) |
| theta | 10242 | -11 | 39 | 23 | 36.80 | Cingulum_Ant_L (9) |
| alpha | 10242 | -22 | 30 | -11 | 30.83 | Frontal_Inf_Orb_L (47) |
| beta | 10242 | -34 | 14 | -34 | 65.48 | Temporal_Pole_Mid_L (38) |
| low gamma | 10242 | 12 | -65 | 58 | 90.62 | Precuneus_R (7) |
| high gamma | 10242 | 12 | -41 | 71 | 53.52 | Postcentral_R (5) |
| DYSLEXIC GROUP |  |  |  |  |  |  |
| delta | 10242 | 40 | -17 | 32 | 18.14 | Postcentral_R (1) |
| theta | 10242 | -10 | -7 | 65 | 45.78 | Supp_Motor_Area_L (6) |
| alpha | 10242 | 8 | 57 | 15 | 36.72 | Frontal_Sup_Medial_R (10) |
| beta | 10242 | -50 | -11 | -21 | 76.89 | Temporal_Mid_L (21) |
| low gamma | 10242 | 18 | -93 | 18 | 120.45 | Occipital_Sup_R (18) |
| high gamma | 10242 | -15 | -65 | 47 | 63.06 | Parietal_Sup_L (7) |
| 2) ISC(con) vs. ISC(dys) |  |  |  |  |  |  |
| delta | 7485 | -27 | -38 | 1 | -6.97 | Hippocampus_L (54) |
| alpha | 98 | 1 | 27 | 10 | -4.41 | Cingulum_Ant_L (24) |
| beta | 6195 | -50 | -14 | -18 | 9.02 | Temporal_Mid_L (21) |
| low gamma | 103 | 44 | -57 | -9 | -5.59 | Temporal_Inf_R (37) |
| high gamma | 834 | 8 | 51 | 20 | -5.46 | Frontal_Sup_Medial_R (10) |

| 3) CORRELATION OF ISCs WITH READING-RELATED MEASURES |  |  |  |  |  |  |
| --- | --- | --- | --- | --- | --- | --- |
| PHONOLOGICAL PROCESSING |  |  |  |  |  |  |
| *delta* | 6451 | -57 | -24 | 26 | 0.24 | SupraMarginal_L (40) |
| *theta* | 1047 | 41 | -63 | 7 | 0.25 | Temporal_Mid_R (19) |
| *alpha* | 91 | -8 | -62 | 48 | 0.15 | Precuneus_L (7) |
| *beta* | 630 | 56 | -15 | 40 | 0.29 | Postcentral_R (1) |
| *high gamma* | 71 | -22 | 44 | 23 | 0.26 | Frontal_Sup_L (10) |
| TECHNICAL READING |  |  |  |  |  |  |
| *delta* | 2395 | -19 | -51 | 2 | 0.18 | Precuneus_L (30) |
| *alpha* | 48 | 5 | 21 | 25 | 0.18 | Cingulum_Ant_R (32) |
| *low gamma* | 3695 | -28 | -70 | -5 | -0.28 | Fusiform_L (19) |
| WORKING MEMORY |  |  |  |  |  |  |
| *delta* | 125 | 14 | 52 | 26 | 0.15 | Frontal_Sup_R (9) |
| IQ |  |  |  |  |  |  |
| *delta* | 1331 | -55 | -23 | 28 | 0.23 | Postcentral_L (1) |

There is an overlap of the ISCs of both groups in all frequency bands, only marginally in the theta band (Supplementary Figure 1). In the delta frequency band, the control participants had significant ISC in temporal, parietal, and central areas; the maximum was in the right mid-cingulate cortex. Dyslexics exhibited ISC in right central and parietal areas, peaking at right postcentral areas. In the theta band, controls had synchronized activity in a defined area depicting the left anterior cingulate cortex, whereas in dyslexics the ISC pattern was more distributed towards left fronto–parietal and temporal areas, and right frontal and temporal areas, peaking at a location roughly corresponding to the left supplementary motor area. In the alpha band, ISC was found in bilateral inferior frontal gyrus, inferior temporal, and frontal areas with peaks in frontal areas in both groups. In the beta band, we observed bilateral frontal and temporal ISCs in both groups and the maxima were in left middle temporal cortex. The low gamma band showed frontal and parietal ISCs in both hemispheres in both groups, and additional strong bilateral occipital ISCs in the dyslexic group only. The high gamma band synchronized in both groups in bilateral superior parietal and postcentral areas that extended into occipital areas in the dyslexic group.

### 3.2 ISC differences between dyslexics and controls

Clusters depicting the brain areas that synchronized significantly differently (corrected for *p* < .05/6) between the control and dyslexic group are shown in Figure 3, and the maximal differences of these areas are summarized in Table 3. The results show that the ISC contrast between the groups manifested in distinct brain areas that differed between frequency bands. Whereas controls synchronized stronger in the delta, alpha, low gamma, and high gamma bands, dyslexics had predominantly stronger ISC in the beta band (Figure 3).

**Figure 3.**
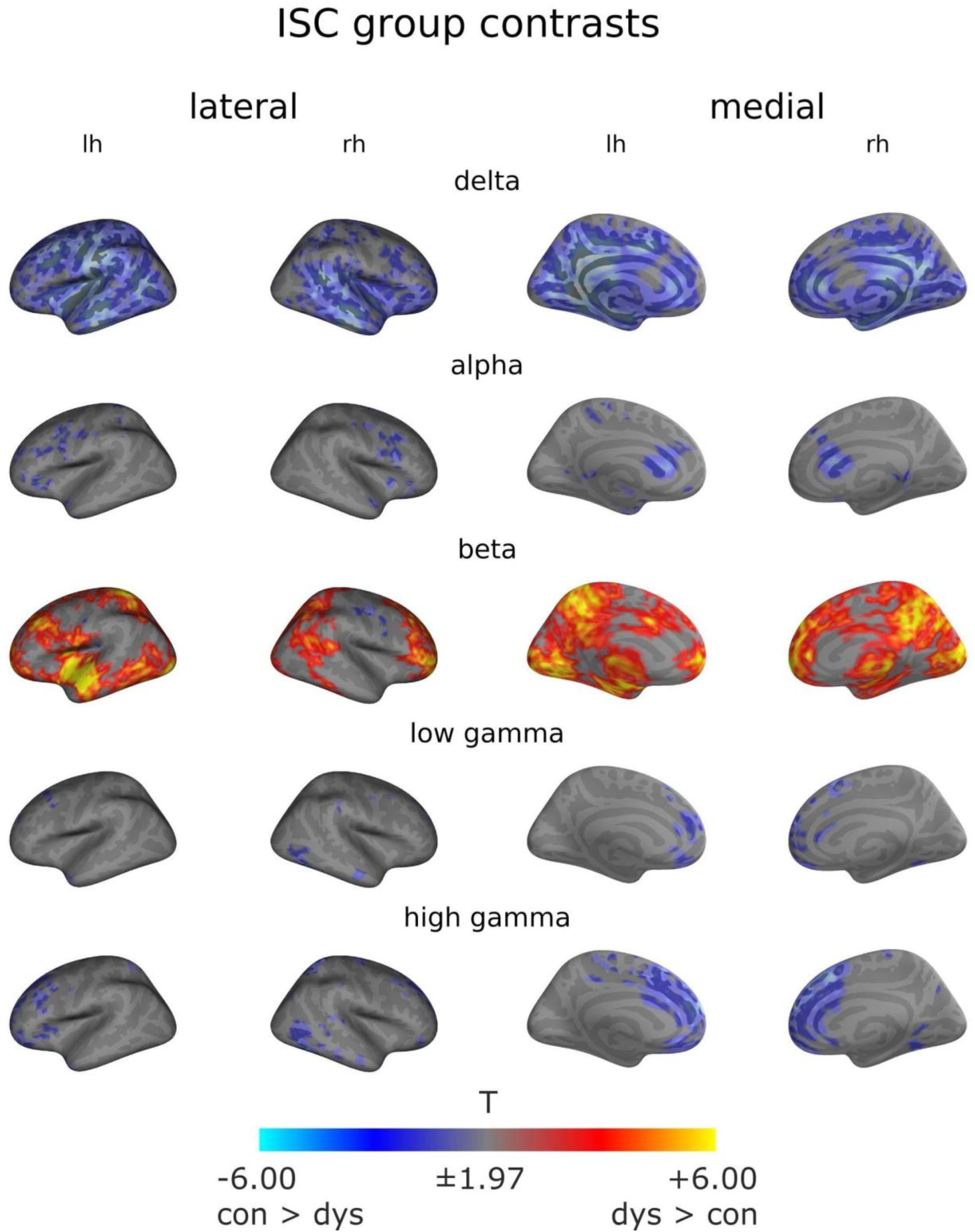
Contrast of inter-subject correlations (ISCs) between the dyslexic and control group for listening to speech. Cold colors indicate stronger ISCs in the control than dyslexic group (con > dys), and warm colors stronger ISCs in the dyslexic than control group (dys > con).

In the delta band, typical readers had significantly stronger ISCs than dyslexics in bilateral auditory cortices, bilateral mid-cingulate cortices, and left central as well as frontal areas. In the theta band, no significant clusters were observed after correcting for multiple comparisons. In the alpha band, frontal and mid-cingulate areas showed higher ISCs in controls than in dyslexics. In the beta band, stronger ISC was found in the dyslexic than control group in a left-hemispheric cluster including superior and middle temporal areas which also contained the maximal difference between the groups, as well as in more focal left-hemispheric occipital pole, superior parietal, and frontal areas. In the right hemisphere, dyslexics synchronized stronger than controls in superior and middle frontal areas including the frontal pole, as well as occipito–parietal areas. Small clusters of enhanced ISC in controls were found in right central areas. In the low gamma band, small clusters of enhanced ISC in controls were found in the right inferior temporal and bilateral frontal areas. In the high gamma band, controls had higher ISC than dyslexics in bilateral frontal, and right temporal areas, peaking in the right superior medial frontal cortex.

### 3.3 Correlation of neuropsychological tests and ISC strengths

The regression matrices showing the mean values of neuropsychological test composites between each subject pair that were used as models for the Mantel test are visualized in Figure 4. All significant correlations of neuropsychological composites and ISCs during listening to speech are visualized as clusters on the *fsaverage* brain in Figures 5 and 6. Only clusters larger than 25 source points were considered significant. Alongside, the mean ISC in the largest cluster was plotted against the neuropsychological composite (for mean ISC vs. neuropsychological composite plots in the second-largest cluster, see Supplementary Figure 2). Significant correlations were found in all frequency bands, being predominantly positive (better reading-related skill was associated with higher ISC), except for technical reading in the low gamma band, where worse technical reading skills were associated with higher ISC in most brain areas The brain areas of the peak correlations between neuropsychological composites and ISC are summarized in Table 3.

**Figure 4.**
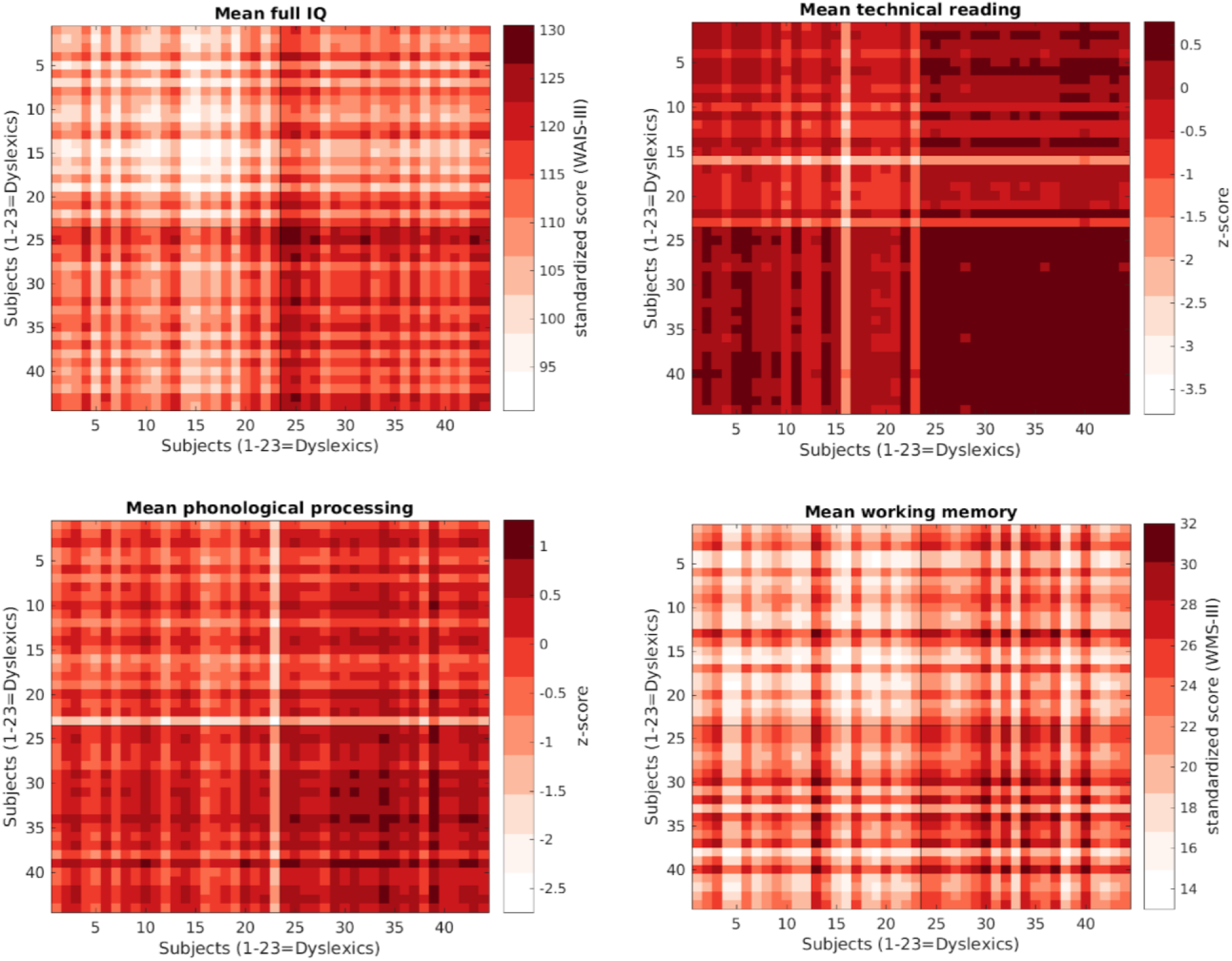
Regression matrices for mean scores of neuropsychological test composites between subject pairs that were used as models for the Mantel test, which tested whether these behavioural models could be explained by the brain ISCs. Z-scores for phonological processing and technical reading. Standardized test scores for IQ and working memory.

**Figure 5.**
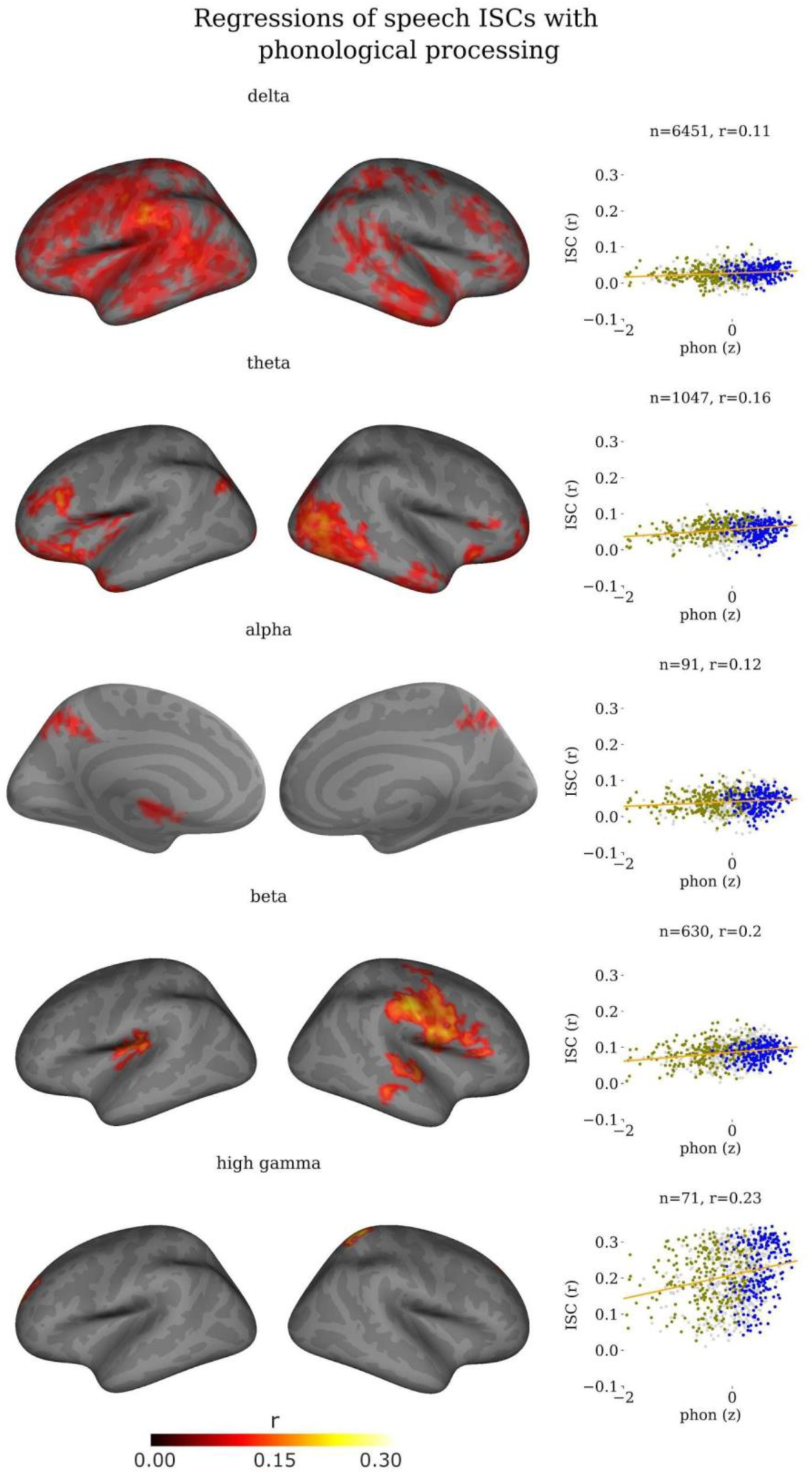
Mantel regressions (r) between phonological processing and inter-subject correlation (ISC) adjusted with cluster correction. Left: Significant regressions on left and right brain hemispheres, lateral views, except for alpha band medial view. Right: Mean ISC (r) in largest cluster plotted against phonological processing score (z) for all subject pairs (ocre - dyslexic pairs, blue - control pairs, grey - mixed pairs) including a linear regression model (orange line). Cluster size (n) and the mean correlation in the largest cluster (r) are indicated above the scatter plots.

**Figure 6.**
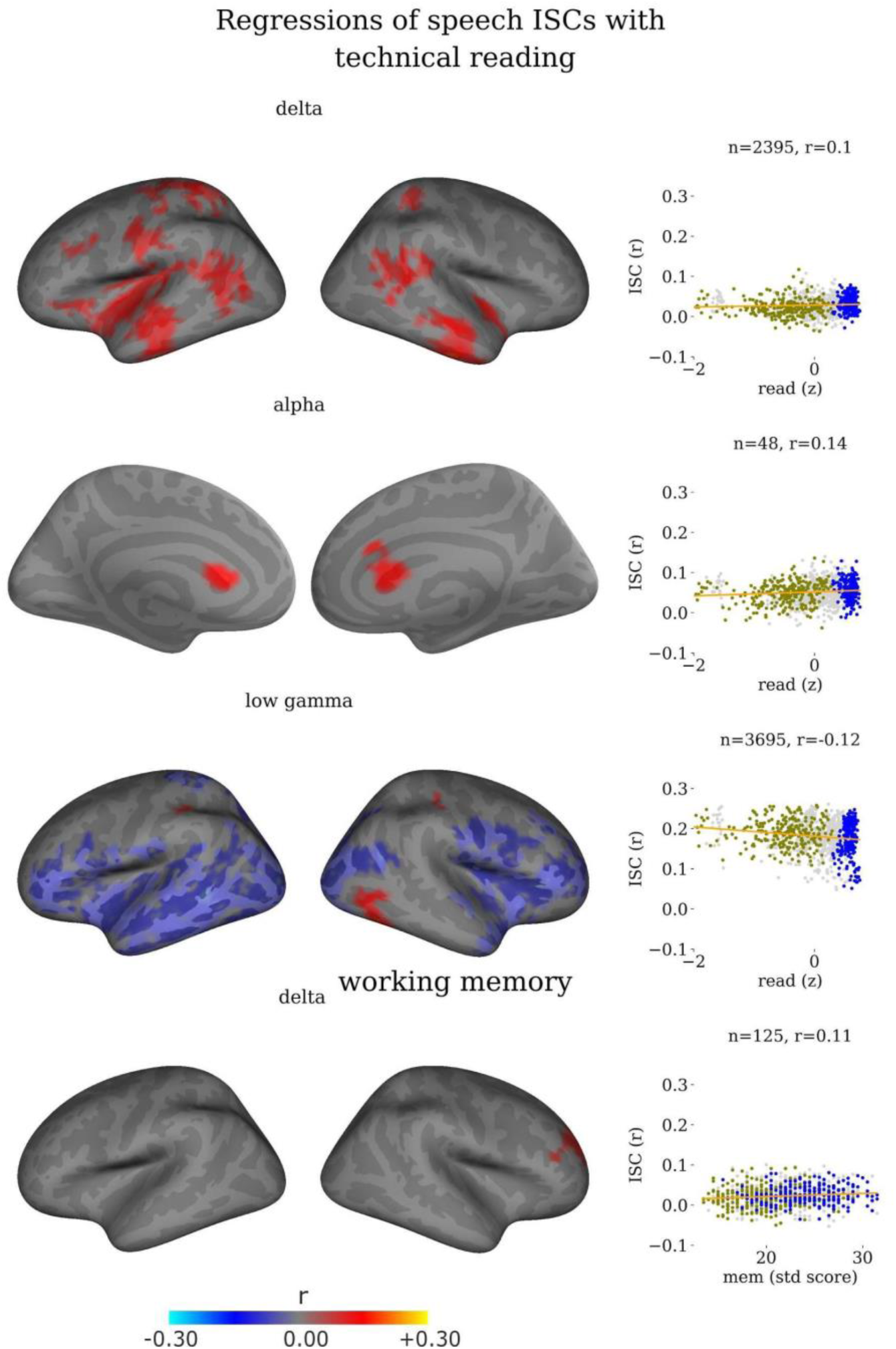
Mantel regressions (r) between technical reading/working memory and inter-subject correlation (ISC) adjusted with cluster correction. Left: Significant regressions on left and right brain hemispheres, lateral views, except for alpha band medial view. Right: Mean ISC (r) in largest cluster plotted against reading score (z) or standardized working memory score for all subject pairs (ocre - dyslexic pairs, blue - control pairs, grey - mixed pairs) including a linear regression model (orange line). Cluster size (n) and the mean correlation in the largest cluster (r) are indicated above the scatter plots.

Phonological processing correlated with ISC during listening to speech in five frequency bands, i.e. all except low gamma (Figure 5). The locations of significant correlations differed between the bands. The largest clusters were found in delta, theta and beta bands. In the delta band, significant correlations were found in left-hemispheric postcentral/superior parietal, precentral, supramarginal, frontal, transverse, middle and superior temporal areas as well as right-hemispheric central, frontal, inferior and middle temporal areas. The maximum correlation in the largest cluster between ISC strength and phonological processing scores was *r* = 0.24 in the left supramarginal gyrus (Table 3). In the theta band, significant clusters were found in left-hemispheric temporal pole, orbitofrontal, rostral middle frontal, and occipital areas. In the right hemisphere, the largest cluster was around the occipital pole extending into middle temporal areas where the peak was located. Other significant correlations were found at smaller inferior temporal and frontal-pole clusters in the right hemisphere. In the alpha band, bilateral superior parietal, and orbitofrontal areas were correlated with phonological processing, showing a maximum correlation at the left precuneus. In the beta band, left-hemispheric insula, and right-hemispheric middle and superior temporal, pre- and postcentral, pars opercularis, pars triangularis, caudal middle and rostral middle frontal areas showed significant correlations between phonological processing and ISC during listening to speech. The maximum correlation was *r* = 0.29 in the right postcentral area. In the high gamma band, small clusters in left superior frontal, and right superior parietal/postcentral areas were significantly correlated to phonological processing skills. The maximum correlation in the left superior frontal cluster was *r* = 0.26.

Technical reading correlated with ISC during listening to speech in the delta, alpha, and low gamma bands (Figure 6). In the delta band, significant regressions between technical reading and ISC during listening to speech were found in the left superior and inferior parietal cortex, central, superior, middle and temporal areas, and insula. Right-hemispheric correlations were located in the inferior and middle temporal cortex, supramarginal, inferior parietal, and postcentral areas. The peak of the largest cluster was at the left precuneus. In the alpha band, bilateral anterior cingulate cortices showed significant correlations with technical reading. Whereas all other regressions indicated that better reading-related skills are associated with higher ISCs, in the low gamma band, also negative associations were found, indicating that worse technical reading was associated with higher ISCs. Negative clusters were found in left temporal and occipital areas, as well as orbitofrontal and superior parietal areas, the largest cluster having a peak at the left fusiform area. In the right hemisphere, occipital and inferior frontal, middle frontal and orbitofrontal areas were negatively associated with technical reading skills. Positive associations were found at a medium-sized cluster in the occipital right hemisphere No significant regressions after corrections were found for the theta, beta, and high gamma band.

Working memory function correlated significantly with ISC in the delta band in a right superior medial frontal brain area (Figure 6). In the other frequency bands, no significant regressions were found.

IQ correlated significantly with ISC in the delta band (Supplementary Figure 3). Left supramarginal, pre- and postcentral, insula, and medial temporal areas showed significant correlations, with the maximum in the left postcentral area. In the right hemisphere, ISCs in medial and inferior temporal areas, rostral middle and lateral orbitofrontal areas, as well as insula, were positively correlated with IQ. In the other frequency bands, no significant correlations emerged.

## 4 Discussion

The aim of the present study was to examine the neural dynamics of dyslexic and typical readers during listening to natural speech. To this end, typical readers and participants with confirmed dyslexia listened to several short excerpts of native Finnish speech while their neural activity was recorded with MEG, which – compared to fMRI – enabled us to analyze the temporal aspect of the neural signal in more detail. We found significant ISC in six commonly investigated frequency bands and could thus delineate neural dynamics at different paces, including the modulations of slow and fast rhythms in the brain. These rhythms are postulated to have neurophysiologically meaningful functions in speech processing (Meyer, 2018).

Firstly, our results confirm and extend the knowledge on between-subjects coupling of brain areas during listening to continuous speech. Secondly, our results suggest atypical ISC patterns during speech processing between dyslexic participants. We found lower ISC between dyslexic compared to typical readers in the delta, alpha, low gamma, and high gamma frequency bands, and mostly enhanced coupling between dyslexics in the beta band. Thirdly, reading-related measures were correlated with the strength of brain-to-brain coupling during listening to speech. The strongest correlations, observed in most of the frequency bands, were found for phonological processing, followed by technical reading, and working memory function.

### 4.1 Interbrain correlation during listening to speech

The ISC patterns we observed in typical readers were overall consistent with those previously found with fMRI during listening to natural speech (Wilson et al., 2008; Stephens et al., 2010; Lerner et al., 2011; Silbert et al., 2014; Finn et al., 2018). These fMRI studies and the results of the present study showed significant ISC in bilateral auditory cortices and language areas along the superior temporal cortex, parietal and midline areas, including precuneus, as well as frontal areas. The present results replicate earlier findings with complex natural stimuli, that is, consistent activation not only in primary sensory cortices but also in higher-order regions (Hasson et al., 2004; Lerner et al., 2011; Finn et al., 2018). Bilateral temporal areas are known to be involved in speech processing and comprehension (see e.g., Hickok and Poeppel, 2007), and therefore were expected to show ISC in our study. In addition, other linguistically relevant and extralinguistic areas showed ISC during listening to speech. Of those, inferior frontal postcentral and parietal areas, specifically premotor areas, belong to a network involved in auditory and speech perception (Giraud and Poeppel, 2012; Schomers & Pulvermüller, 2016; Lima et al., 2016). Moreover, precuneus has been shown to play a role in higher-level social processes, such as role or perspective taking and episodic memory retrieval (Cavanna & Trimble, 2006), and it was suggested to be part of the theory-of-mind network together with STS and temporal-pole areas (Mar, 2011).

In addition, our dyslexic participants displayed ISC in occipital areas, for which previous fMRI studies have not reported ISC during listening to speech. Synchronized activity in occipital areas has recently been shown to support mental imagery and the elicitation of individual meanings of a narrative (Saalasti et al., 2019).

ISC in the beta band was maximal in the left temporal pole in the control group. Temporal pole has been previously associated to speech processing (Tzourio et al., 1998) as well as to semantic word processing or perception (Crinion et al., 2006; Marinkovic et al., 2003) and memory retrieval (Fink et al., 1996). Also the functional role of the beta band was suggested to be lexical–semantic prediction during speech comprehension (Lewis et al., 2015; 2016). Therefore, our results of maximal beta-band ISC in the left temporal pole could reflect processing of meanings of words in the continuous speech.

### 4.2 ISC differences between dyslexics and controls

To assess whether the extent of ISC differed between the dyslexic and control group, we compared the pairwise correlation maps between the two groups. We found that ISC was different between the groups in all frequency bands, however, with different patterns across the frequency bands. In the delta, alpha, low gamma, and high gamma bands, typical readers showed enhanced ISCs compared to dyslexic readers. On the other hand, ISC was mostly stronger in dyslexic than typical readers in the beta band.

The enhanced ISC in the delta band in typical readers compared to dyslexics is consistent with the temporal sampling deficit theory (Goswami, 2011), which predicts that dyslexics especially in lower frequency bands would show a reduced sampling of information contained in the continuous speech stream. Delta-band synchronization is thought to be involved in the segmentation of intonation phrases (Giraud and Poeppel, 2012; Meyer, 2018). A reduced brain-to-brain coupling in this frequency band could therefore be indicative of deficits in temporally synchronized sampling of phrase boundaries. Previously shown reduced neural entrainment to the speech envelope in the delta band in dyslexics compared to typical readers (Molinaro et al., 2016) corroborates our results. Also phase locking to speech modulations at the delta rate was found to be atypical in dyslexia (Hämäläinen et al., 2012), suggesting additional delta-rate speech processing deficits.

Alpha-band ISC was diminished in dyslexic compared to typical readers in bilateral frontal and mid-cingulate areas. The syllabic rate in speech has its upper limit in the alpha range (around 10 Hz). A less synchronized brain activity in the alpha band could therefore imply inaccurate parsing of syllables in dyslexia. Syllable tracking has often been suggested to occur mainly in the theta band (Luo and Poeppel, 2007; Meyer 2018). However, we did not observe less (or more) synchronized theta activity in dyslexics. Similarly, De Vos and colleagues (2017a) did not find atypical brain responses to theta-band speech modulations in dyslexic adolescents, but less synchronized activity to alpha-rate modulations. Alternatively, alpha oscillations have been suggested to aid the storage of syntactic phrases (Meyer 2018). The less synchronized alpha-band activity could suggest an inefficient storage of syntactic phrases to memory.

The enhanced beta-band ISCs in the dyslexics compared to controls support our hypothesis of enhanced coupling in higher frequency bands in dyslexia. De Vos and colleagues (2017b) showed that dyslexic children – when beginning to read – exhibited larger auditory steady-state responses to speech-weighted noise amplitude-modulated around 20 Hz (beta band), referred to as phoneme-rate modulations by the authors. This higher neural synchronization to phoneme-rate modulations was correlated with poorer reading and phonological skills in that study. Similarly trending results were obtained for dyslexic adolescents (De Vos et al., 2017a). In that light, our findings support the ‘oversampling’ hypothesis brought forward by Lehongre and co-workers (2011). According to this hypothesis, phoneme-rate information reflected in the beta band could be oversampled, resulting in working-memory overload and therefore slower or less accurate extraction of phonemic information from speech. Alternatively, enhanced synchronization in the beta band has been suggested to be a compensatory mechanism for the processing of phonemic-rate information (De Vos et al., 2017a). The maximal ISC difference in the largest cluster between the groups localized in the left middle temporal cortex. In terms of phoneme processing, the left middle temporal cortex would be expected to play a major role, as it is an integral part of speech and word processing (Hickok and Poeppel, 2007). In fMRI studies, the peak location for differences between our groups found for the beta band has been frequently associated with activations during listening to speech in various ways (Narain et al., 2003; Oechslin et al., 2010; Straube et al., 2013b; Nagels et al., 2013; Evans et al., 2016, Wolf et al., 2017).

In both low and high gamma bands, we found weaker ISCs in dyslexic readers than in controls. This result was rather unexpected, as we hypothesized that in higher frequency bands dyslexics could show enhanced ISCs than controls (Lehongre et al., 2011). The gamma band as a whole (usually > 30 Hz) has been associated with numerous functions in speech processing, such as phonemic processing (Giraud and Poeppel, 2012), long-term memory processing (Ward, 2003), lexico–semantic retrieval (Pulvermüller et al., 1996; Mai et al., 2016) as well as tracking of phrase and syllable rhythms in continuous speech (Ding et al., 2015).

Low-gamma-band group differences with weaker ISCs in dyslexics compared to controls were found in a few small clusters, the largest one being in right inferior temporal areas. Activation differences at this location have been reported between dyslexics and controls in fMRI studies with visual and phonological tasks (Eden and Zeffiro, 1998; Peyrin et al., 2012). Using MEG, Lehongre and colleagues (2011) found impaired left-hemispheric sensitivity in temporal areas in dyslexia to sounds that were modulated in the range of the low gamma band. While their and our results agree on a weaker sensitivity or ISC, the source locations differ. Our results suggest weaker ISC in the right temporal areas and in bilateral medial frontal areas in dyslexics. These differences could be due to different stimuli (sound modulations vs. natural speech) and analysis methods (frequency tagging vs. ISC).

The natural stimulus presentation in the present study differs from the well-controlled designs often used in event-related neurophysiological studies. Despite the different paradigms, event-related brain responses are commonly filtered in the range of from delta to beta or low gamma frequencies (i.e. around 0.5 to 30 Hz), and therefore the evoked-response-based findings on dyslexia (for reviews, see Hämäläinen et al., 2013; Kujala and Näätänen, 2001) may aid the interpretation of our ISC results. Sources of these responses during language-related tasks suggest functional differences between dyslexic and typical readers in left and right perisylvian language regions (for a review, see Heim and Keil, 2004). The results of the present study may reflect certain brain synchronization patterns that occur due to salient events in the continuous speech. As discussed in more detail above, these events may be related to different hierarchies of speech, such as phonemes, syllables, phrase boundaries etc. We found maximal ISC differences in the largest clusters between typical and dyslexic readers in this study in left and right perisylvian language areas in the beta and low gamma band. In the beta band, dyslexics had stronger ISCs in the left middle temporal cortex than typical readers, and in the low gamma band, typical readers had stronger ISCs in the right inferior temporal cortex. Stronger ISC in the left temporal cortex in dyslexics could be seen as a contrast to previous results, such as weaker evoked-response sources in left temporo-parietal regions in children with dyslexia during reading (Simos et al., 2000a; Simos et al., 2000b) or atypical functional organization of the left auditory cortex in dyslexia during pure-tone and syllable processing (Heim et al., 2000). Weaker ISC in the right temporal cortex in dyslexics is supported by findings of altered source configuration in the right hemisphere in dyslexia during processing of repeated syllables (Heim et al., 2004). While both these studies and our study investigated speech or related processes, the differences in stimuli and analysis raise the question whether they are comparable.

Most of the above-mentioned studies that investigated oscillations during speech processing have looked at how brain signals in different frequency bands were following the speech signal. However, inter-subject synchronization during processing of speech has been studied to a much smaller extent. Our results show for the first time with MEG the synchronous neural processes between participants during speech processing, complementing earlier studies that investigated brain-to-stimulus coupling. The current approach focuses on how similarly speech was processed in the target groups, and how the synchronous neural processes differ between participants with and without dyslexia.

### 4.3 Correlation of neuropsychological tests and ISC strengths

ISC of both groups was significantly correlated with the neuropsychological composites of phonological processing, technical reading, and working memory. Correlations were found in most frequency bands for the phonological processing composite, followed by technical reading and working memory.

The phonological processing composite consisted of the ‘Pig Latin’ test, non-word span length, digit span length, and rapid alternating stimulus naming, all tapping into processing of phonological information. Large brain areas in delta, theta, and beta bands were positively correlated with phonological processing across both groups, meaning the stronger the brains synchronized, the better phonological skills the subjects had. A maximum correlation in the delta band was found in the supramarginal gyrus which incidentally was also the only area consistently correlated with IQ differences. The association between dyslexia and IQ has been a topic of debate for many years now (e.g. Shaywitz et al., 1995; for a review, see Stuebing et al., 2002). Following the recommendation of Dennis and colleagues (2009), we did not use IQ as a covariate, but rather investigated its association with ISC separately. In the theta band, the largest cluster indicating significant correlations could be located in the right middle temporal and occipital areas: higher ISC was associated with better phonological processing skills. Therefore, it could be that increased ISC in those areas reflects better speech parsing, thus leading to better phonological skills. In the beta band, the ISC in a large cluster around the right postcentral area was associated with phonological processing skills. According to the direct group comparison, this area was more strongly synchronized in typical than dyslexic readers, although in many other areas the opposite contrast was observed. It is possible that the phoneme information, the parsing of which is reflected in the beta band (De Vos et al., 2017b), was processed inefficiently by dyslexic readers in the postcentral right-hemispheric area and therefore the lower ISC was associated with worse phonological processing skills. In other words, typical readers with better phonological processing skills could be more efficient in processing phonemes reflected by higher ISC. Less entrainment to acoustic modulations around 30 Hz in dyslexics has also previously been associated with worse phonological processing, but better rapid naming skills (Lehongre et al., 2011). Due to the use of different subtests for phonological processing (the phonological processing composite in our study contained rapid naming as one of the subtests whereas Lehongre and colleagues (2011) separated phonological processing and rapid naming) and slightly different frequency limits (upper limit for the beta band was 25 Hz in our study) it is unclear whether their and our results tap on the same processes.

The technical reading composite comprised word and pseudoword list reading scores in speed and accuracy. Thus, this score merely reflects reading skills at the single-word level, but not, e.g., reading comprehension. Technical reading was positively associated to the ISC strength during listening to natural speech in the delta band, with the largest cluster at the left precuneus, a higher correlation between participants reflecting better technical reading scores. Although some of the brain areas that were correlated with technical reading overlap with those that correlated with IQ, the maxima differ. In line with the group differences in the delta band, a lower correlation between dyslexic participants is associated with worse technical reading skills. Low-level auditory processing could be related to the processing of phrase boundaries, corresponding to the delta-band frequencies (Giraud and Poeppel, 2012; Meyer, 2018). Abnormal low-level auditory processing can lead to impaired speech representations in the brain, which can affect reading abilities as in dyslexia (Bailey and Snowling, 2002; Goswami, 2015). In the low gamma band, the largest ISC cluster showed negative correlations with technical reading skills. Left temporal areas were included in this largest cluster, whereas right temporal areas did not show significant correlations, except in a small cluster of positive correlations. As the metric of technical reading skills is saturated in controls, it is possible that a higher ISC in left temporal areas in dyslexics reflects a compensatory mechanism for phoneme processing. Even though we found negative correlations between ISC and technical reading in the left hemisphere, in the group comparison, these areas did not have higher ISC in dyslexics. The small cluster of positive associations between technical reading and ISC, on the other hand, corresponded to the same area with stronger ISC in controls in the group contrast.

Working-memory capacity correlated with ISC strength only in the delta band. The correlation in such a low frequency band was rather unexpected as Lehongre and colleagues (2011) previously associated a working-memory deficit with enhanced entrainment to rates above 40 Hz, i.e., in the higher gamma range. The right superior frontal area that was maximally correlated with working-memory capacity in the delta band did not appear to be significantly different between groups, although the direction of correlation suggests that a higher ISC would be associated with better working-memory skills, and these skills in our two groups are significantly different from each other. Associations with the delta-band have not been reported before and could be looked at in follow-up studies employing different methods. Possibly, a within-group correlation analysis could reveal further directions.

### 4.4 Limitations and future directions

The interpretation of ISC is the first limitation we want to address. First, for a certain brain region, ISCs in two frequency bands may also be explained by cross-frequency coupling (Canolty et al., 2010; Giraud & Poeppel, 2012). The ISC method used in this study is not adequate to disentangle cross-frequency coupling from independent synchronization in multiple frequency bands, and it should be investigated in *a-priori* defined bands and regions of interest, if applicable, with different methods, using both phase and amplitude information.

Future studies could investigate the effect of the age of the participants. Our participants were adults, and therefore the ones with dyslexia may have employed different compensation mechanisms and strategies for reading, which should be reflected as differences in those brain processes that are synchronized. A natural follow-up of this study would be to investigate these processes in children of different ages, i.e. before and after reading acquisition, to determine whether the atypical synchronization effects in dyslexia are rather due to genetic or environmental influences.

Another important point is the interpretation of cluster-based permutation tests. One should be aware that the results of these tests do not return a real spatial extent of the “significant” clusters (Sassenhagen & Draschkow, 2019). Therefore, the obtained shapes of the significant clusters are only observational. Despite those limitations, the cluster-based permutation tests are powerful in controlling for multiple comparisons in the high-dimensional MEG ISC matrices and were therefore the method of choice.

## 5 Summary and conclusions

With our novel approach of frequency-band-specific inter-subject correlation of MEG acquired during listening to natural speech, we showed that the strength of ISC differs between dyslexic and typical readers, with weaker ISCs in dyslexics in the delta, alpha, low and high gamma bands, and stronger ISC in dyslexics in the beta band. Furthermore, the strength of ISC was associated with phonological skills as well as technical reading and working-memory function. Our findings shed light on how speech processing is reflected in different MEG frequency bands in healthy adults and in those with reading impairments and suggest how these brain dynamics are associated with behavioural outcomes. Unveiling speech processing in the brain in ecologically valid conditions can help uncover the complex neural basis of dyslexia.

## Supporting information

speech_transcription_eng

speech_transcription_fin

## Acknowledgements

The authors thank all research assistants involved in the MEG and MRI recordings and neuropsychological testing. We would like to thank Emma Suppanen for providing us with the original version of the Matlab ISC scripts. Finally, we would like to thank all participants for their engagement in this study. This work was supported by Jane and Aatos Erkko Foundation and the Academy of Finland [project numbers 276414 and 316970]. The first author received funding from the Research Foundation of the University of Helsinki.

