## Supplementary Figures for "Atypical MEG inter-subject correlation during listening to continuous natural speech in dyslexia"

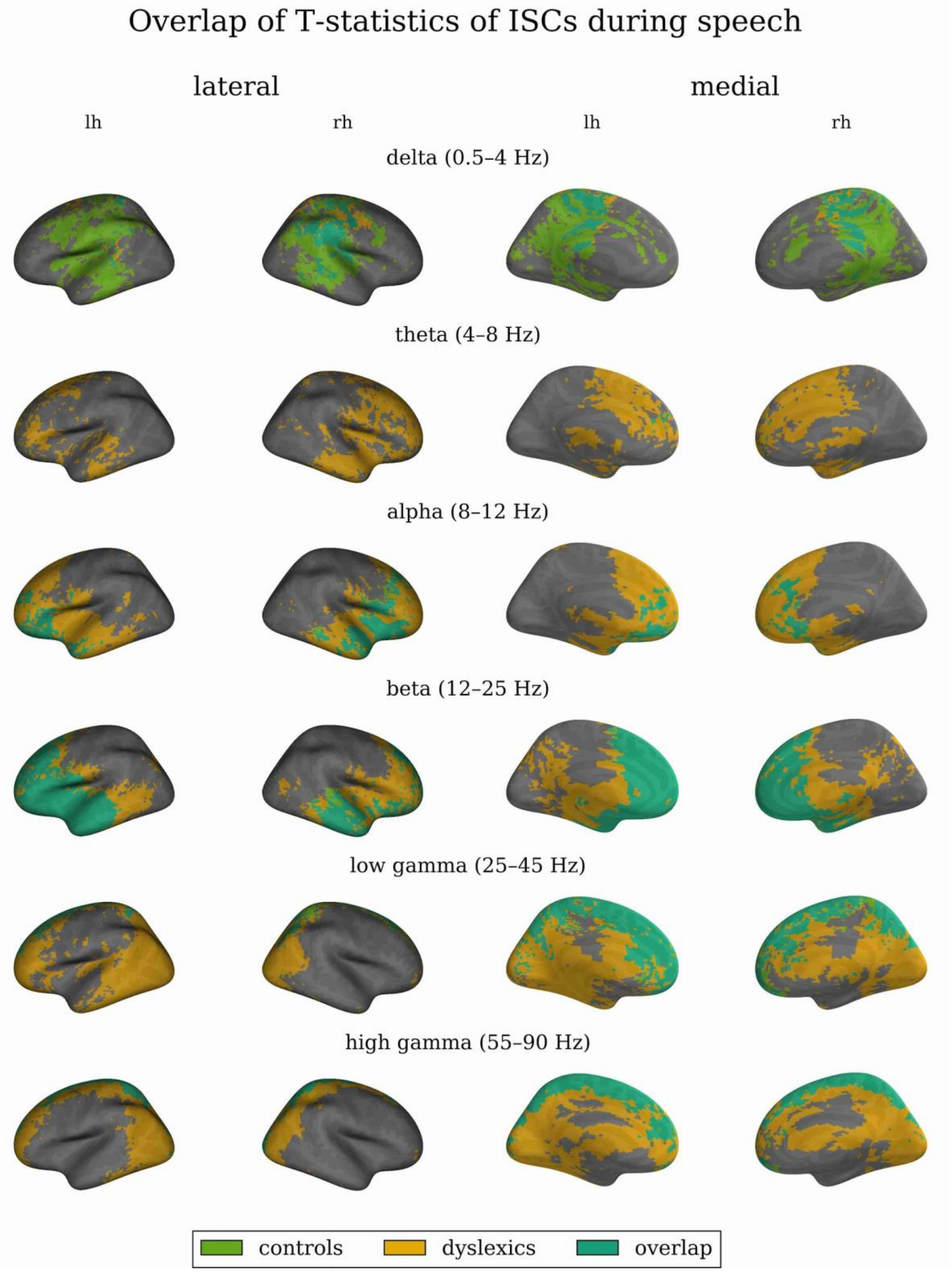


*Supplementary Figure 1. Overlap of T-statistics of permutation-based one-sample t-tests for inter-subject correlations (ISCs) during listening to speech in control and dyslexic group. Only ISCs above the cutoff T-values as chosen in Figure 2 (lowest values in the T-legends) are visualized, separately for the control group only (green), dyslexic group only (yellow), and the overlap of both groups (turquoise). Lateral (first two views) and medial (last two views) views (lh – left hemisphere, rh – right hemisphere) are shown for each frequency band.*

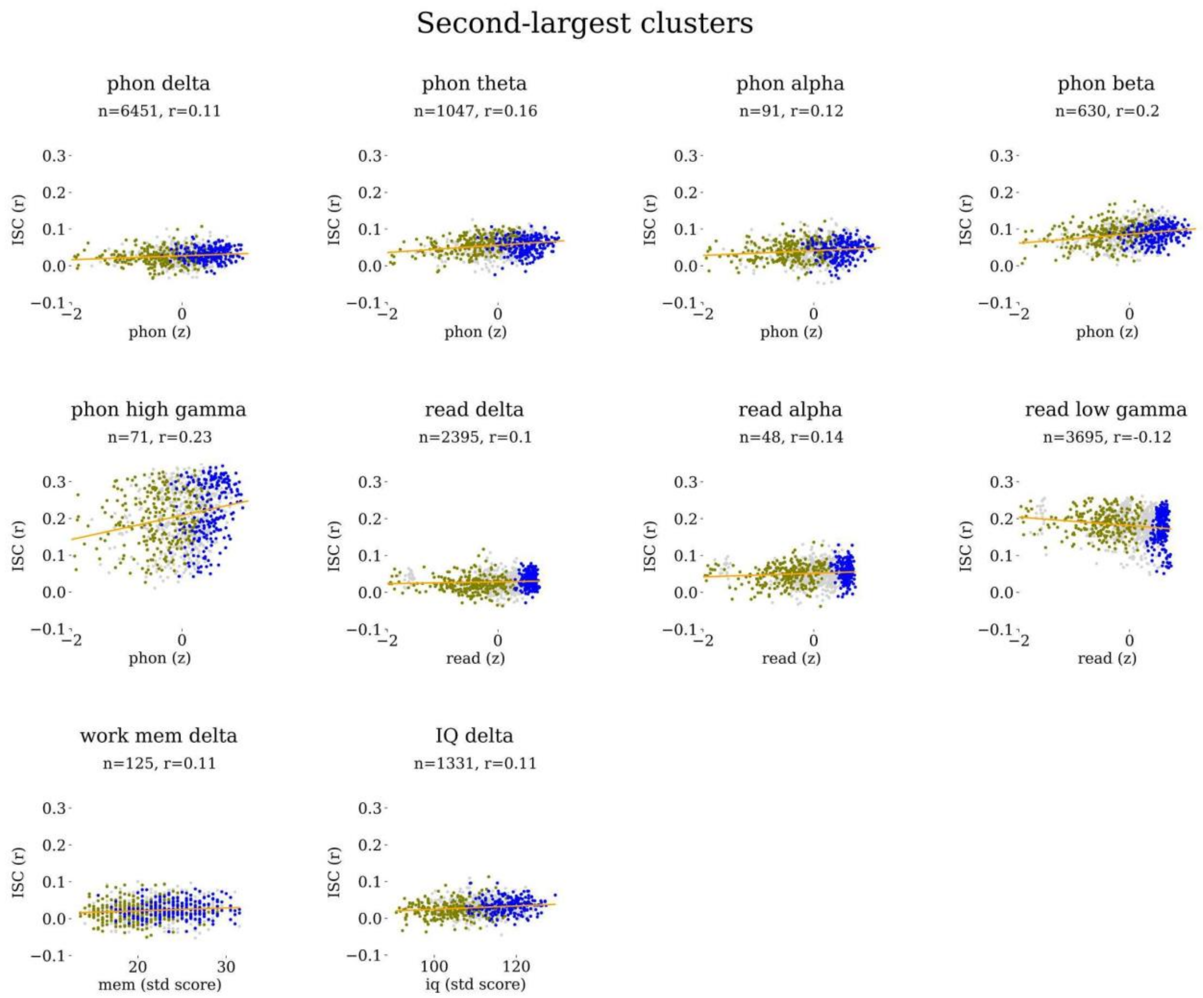


*Supplementary Figure 2. Mean inter-subject correlations (ISC, r) in second-largest cluster plotted against phonological processing score (z), technical reading score (z), standardized working memory score, or standardized IQ score for all subject pairs (ocre - dyslexic pairs, blue - control pairs, grey - mixed pairs) including linear regression line (orange). Cluster size (n) and the mean correlation in the second-largest cluster (r) are indicated above the scatter plots. Only combinations of reading-related skill and frequency bands (delta, theta, alpha, beta, low gamma, high gamma) that had significant second-largest clusters are depicted. phon - phonological processing, read - technical reading, work mem - working memory, IQ - intelligence quotient*

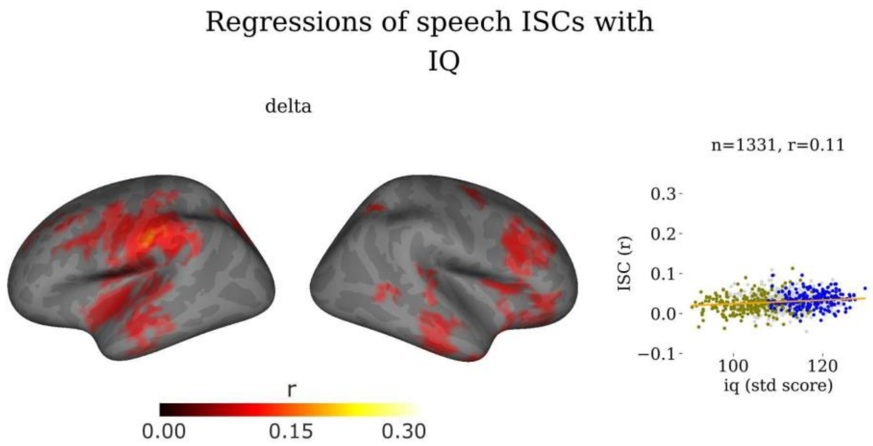


*Supplementary Figure 3. Mantel regression (r) between IQ and inter-subject correlation (ISC) adjusted with cluster correction. Left: Significant regressions on left and right brain hemispheres, lateral views. Right: Mean ISC (r) in largest cluster plotted against IQ score (standardized score) for all subject pairs (ocre - dyslexic pairs, blue - control pairs, grey - mixed pairs) a including linear regression model (orange line). Cluster size (n) and the mean correlation in the largest cluster (r) are indicated above the scatter plot.*
