## Supplementary material for "Atypical MEG inter-subject correlation during listening to continuous natural speech in dyslexia": speech_transcription_eng

---------------------------------------------------------------------------------------------------------------------------------------

This free speech was used as stimulus in **Thiede et al. Atypical MEG inter-subject correlation during listening to continuous natural speech in dyslexia**

<https://doi.org/10.1101/677674>

**Please cite this paper, if you use the stimulus or any derivation thereof!**

---------------------------------------------------------------------------------------------------------------------------------------

**Part 1**

Hi! Excuse me, can I bother you for a minute?

Yes, of course.

Yeah, we are apparently in Kamppi Centre, I’m a little lost.

Yes, this is Kamppi Centre, where are you headed?

Yeah, I need to find my way to Esplanade Park.

Oh, okay, well... How would it now be easiest for you... Well maybe so that if this is the main door to Kamppi center, you will exit it to a large square. Then you should walk cross this square and well, then comes a big road. Do you know Mannerheimintie?

Yes, yes, I think I just came from there.

Okay, well, there at Mannerheimintie you go right, looking from here.

Okay, okay.

And then after a while, you need to walk a couple of blocks, then after a while you will see on the left, there is, do you know this Swedish Theater, it is this big round building?

It might be that I recognise it.

Okay.

Yeah.

So, well, it comes across you on the right and then there behind the theater is Esplanade Park.

Okay, so, back to Mannerheimintie, then right and then a while straight until Esplanade Park on my left.

Right.

Okay, well.

Good, I think you will find it.

Brilliant, thank you!

---------------------------------------------------------------------------------------------------------------------------------------

Yle News at 9 am with Petteri Löppänen, good morning! Employee union’s protest hinders the traffic in the whole country. Busses have been standing still for a day after midnight, trains came to a halt on early morning and rails remain empty until 6 pm. In the capital region metro and trams are missing from the tracks also, and some flights have been cancelled as well. In advance protest was thought to increase traffic jams especially in capital region, but also in other cities across the country. In capital region the morning traffic has flowed quite smoothly and the largest amount of traffic so far has been observed already before 7 am, tells chief administrator officer Mika Jaatinen from Helsinki Traffic Center. “The traffic has been quite all right this morning, the traffic has started to increase about an hour earlier than during normal mornings. That is, in that sense people have followed media well about those possible traffic jams. But, indeed, the traffic has increased maybe an hour earlier compared to normal mornings, at this moment there is a little less traffic on the roads than on normal mornings. So right now we would normally have the rush hour peak of the morning. But right now there is less traffic than during the normal mornings.” The Traffic Center does not comment on afternoon traffic yet, since the morning traffic is still ongoing. The air traffic at Helsinki-Vantaa airport has had only minor problems. Finavia, the operator responsible for airport functions, tells that next moment of suspense will be at the time of the actual protest from 11 am to 1 pm. During that time work is tried to be done by foremen.

---------------------------------------------------------------------------------------------------------------------------------------

Last week I talked about how life does not obey our plans. However, it doesn’t mean that I wouldn’t believe in my dreams and making them to come true. Quite the opposite. Many of my great dreams have come true and they give me great joy even today, daily. On my list of fulfilled dreams over the past years are, among the other things, a big change at work, quitting smoking, learning how to drive a motorcycle and exercising myself from a lousy shape couch potato to a better than average fit adult. My intention is not to brag. Making dreams come true is not about magic dust sprinkled by fairies or exceptional willpower. I’m deep down a comfort-seeking human being. I have just by chance found a working recipe that I have kept repeating from one dream to the next. Because it is not a secret, I will tell it to you. Everything starts with dreaming. You have to dream big and broad. Dream about things that might at first seem unreachable. When a wish keeps coming to my thoughts day after day, week after week and month after month, I understand, I need to reach for it. This incubating period can be long, even years. Well dreamed is halfway done. The mind has an immense power. When I can imagine my dream come true, I know that it is possible also in reality. A dream must scare you, tempt you and excite you at the same time. When you think of it coming true, you feel a pinch in your stomach. A real and own dream is recognised by the big feelings it invokes in you. A dream never feels insignificant.

---------------------------------------------------------------------------------------------------------------------------------------

**Part 2**

What kind of flat, tell me more about it…

Yeah, so, I have this kind of shared two-room flat, 62 square meters there, it is so nice. Only that we are now in fact moving out with my roommate, so that will be in the end of next month, I‘ll be moving to the other side of the park . There is this nice one-room apartment reserved for me. But I’ll be staying in Arabianranta, since I’m so enchanted by it. But yeah well, moving out ahead.

Is your roommate coming to live with you?

No, she’s not, she is actually moving in together with her boyfriend.

Okay.

So I’m moving all alone. But it is great, because I have lived some many years with these roommates. So it is quite nice to try my own wings and see how the life will turn out to be in one-room apartment.

Well, I believe it will turn out quite alright.

Mmh.

It will be like nothing.

Have you lived in shared flats or one-room apartments?

Yes, I do live actually at the moment, it is this kind of shared flat.

Okay.

So that there is kind of two similar rooms and then shared kitchen and the rest, but my roommate is my friend, so it has worked out very well.

Mmh, it is really a very good solution.

Yes, and then we are both studying the same subject, so that we can always wrap our heads around these exercises at the same… at the same time.

Mmh.

---------------------------------------------------------------------------------------------------------------------------------------

It took me a long time to understand where he came from. Little Prince, who himself kept asking me a lot, seemed never to hear my questions. Coincidentally uttered words revealed me this little by little. So he asked, after seeing my aeroplane (I am not drawing my aeroplane, it would be a task too difficult for me):

“What a gadget is that?”

“It is not a gadget, it flies. It is an aeroplane. It is my plane.”

I was very proud being able to tell him that I fly.

To that he proclaimed: “Is that really so! So you have fallen down from sky.”

“Yes”, I replied modestly.

“Oh, that is funny”, and Little Prince burst into joyful laughter, which irritated me very much. I had expected that he would adopt a more serious approach to my accident. Then he added:

“So you too are coming from the sky, which star are you from?”

I began to see a little, how his presence here could be explained and I asked him quickly:

“So you come from a different star?”

But he didn’t reply me. Looking at my plane he shook his head.

“It is clear, you couldn’t arrive from afar with that kind of gadget”. And he fell into deep thoughts.

Then he pulled my sheep out of his pocket and was absorbed observing his treasure. You can imagine, how this hint about other stars had excited my curiosity. I tried to get more clarification on the matter.

“Where are you coming from little man? Where is your home? Where do you want to take my sheep?”

After a quiet pondering he replied: “It was good of you to give me the box. It can well be the house for the sheep during the night.”

“Yes it can and if you are nice, I will give you a cord and a pole, so that you can tie the sheep during the day.”

My suggestion seemed to somehow puzzle the Little Prince.

“Tie it down. What a funny thought.”

“But if you don’t tie it, it can go anywhere and get lost.”

My friend started laughing again.

“Where could it go?”

“Anywhere. Straight ahead...”

Then Little Prince noted seriously: “It doesn’t mean a thing, on my star everything is so small.” And I think he added little sad: “You can not go straight ahead very long.”

---------------------------------------------------------------------------------------------------------------------------------------

Hi! Paula hello!

Hi!

You were then at Summer in Prague, were you not?

Yes, I was.

Was it a good trip?

Well, yes it was, yeah that’s right we haven’t talked about it.

Yes.

Well yes. We were there for a week and it was a sufficient time to… to see all the main things, I think.

Yes. How did you like the city in general?

Well, the city is, like just walking there is maybe the best thing.

Hmm

It's really charming historically and it's just those bridges and it's river and ... Or I think that that is like the best thing in Prague, just looking around and…

Yeah, it is a rather old city, isn’t it?

So yes, yes it is. And just like this central European historic city, where there is.. a river going through and there are beautiful historical bridges and then there is some big cathedral and and…

Yeah.

Yeah, just like that.

Yeah. I was also in fact in Prague a year ago…

Oh, really!

And it was somewhat confusing that the roads are not directly mapped, like.

Oh, you say it! I was lost so many times in there! All the roads in there go according to those roundabouts, like big circles.

Yeah, yeah! It was incomprehensible. I had just told my friend during that rip, that I’m so terribly good at reading a map…

Oh no!

And I kept taking us all the time to completely wrong streets. And I felt like those streets were moving at that time…

Yeah.

..when we went onwards.

Yeah, yeah, I had quite the same feeling and my boyfriend mocked me that you can’t find your way at all.

Yeah, yeah. But is a very nice place.

---------------------------------------------------------------------------------------------------------------------------------------
