## Supplementary material for "Atypical MEG inter-subject correlation during listening to continuous natural speech in dyslexia": speech_transcription_fin

---------------------------------------------------------------------------------------------------------------------------------------

This free speech was used as stimulus in **Thiede et al. Atypical MEG inter-subject correlation during listening to continuous natural speech in dyslexia**

<https://doi.org/10.1101/677674>

**Please cite this paper, if you use the stimulus or any derivation thereof!**

---------------------------------------------------------------------------------------------------------------------------------------

**Part 1**

Moikka! Anteeks saaks mä vähän häiritä?

Joo totta kai.

Joo, me ollaan ilmeisesti nyt Kampin keskuksessa, mä oon vähän eksyksissä.

Joo, tää on Kampin keskus, mihin sun pitäis päästä?

Joo, mun pitäis löytää tonne Esplanadin puistoon.

Aa okei no niin no tota. Mitenköhän nyt sun olis kaikista helpoin. No ehkä silleen että jos tässä on nyt tää Kampin keskuksen pääovi niin niin sä tuut sellaiselle isolle aukiolle. Niin pitäis kävellä sen aukion poikki ja tota sitten tulee se iso tie. Tiiäksä Mannerheimintien?

Joo joo, sielt mä tulin äsken.

Okei no ni, niin sit sieltä Mannerheimintielle lähet sinne oikeelle päin, täältä katsoen.

Okei, okei

Ja sit vähän ajan päästä, sun pitää kävellä pari korttelia, ni sit vähän ajan päästä vasemmalla lähtee sitten, siin on, tiiäksä sellaisen ruotsalaisen teatterin, se on semmoinen iso pyöree rakennus?

Saattaa olla et tunnistan.

Joo

okei

Joo niin se tulee siinä oikeella vastaan ja sit siitä teatterin takaa lähtee sitte Esplanadi

Okei, elikkä, takaisin Mannerheimintielle, sitten oikeelle ja sitten vähän aikaa eteenpäin, kunnes vasemmalla on Esplanadi.

Just niin.

Noni.

Hyvä, etköhän sä löydä.

Loistavaa, kiitos sulle!

---------------------------------------------------------------------------------------------------------------------------------------

Yle uutiset kello yhdeksän ja Petteri Löppänen hyvää huomenta. Työntekijäjärjestöjen mielenilmaus vaikeuttaa liikkumista koko maassa. Linja-autot pysähtyivät pääosin vuorokaudeksi aamuyöllä, junaliikenne seisahtui aamutuimaan ja raiteet pysyvät tyhjinä kello kahdeksaantoista asti. Pääkaupunkiseudulla raiteilta puuttuvat myös metrot ja raitiovaunut, myös lentoja on peruttu. Mielenilmauksen arveltiin ennakkoon lisäävän ruuhkia liikenteessä varsinkin pääkaupunkiseudulla, mutta myös kaupungeissa muualla maassa. Pääkaupunkiseudulla aamuliikenne on sujunut kuitenkin melko jouhevasti ja suurimmat liikennemäärät on toistaiseksi nähty jo ennen seitsemää, kertoo liikennekeskuspäällikkö Mika Jaatinen Helsingin liikennekeskuksesta. ”Liikenne on sujunut ihan hyvin tässä aamusta, elikkä liikennemäärät on lähtenyt ehkä noin tuntia aikaisemmin kasvuun kuin mitä normi aamussa. Eli siinä mielessä ihmiset on hyvin seurannut mediaa noitten mahdollisten ruuhkien osalta. Mut tosiaan liikennemäärät lähteneet aamusta tuntia aikasemmin kuin normaalisti kasvuun, tällä hetkellä liikennettä on jonkun verran vähemmän kuin mitä normaaliaamussa olisi. Et nythän me elettäis normaalia aamuruuhkan huippuhetkiä. Mut nyt liikennettä on siis vähemmän kuin normaaliaamuna.” Iltapäivän liikennettä liikennekeskuksessa ei haluta vielä arvioida, koska aamuliikenne on vielä kesken. Helsinki-Vantaan lentoliikenteeseen on aiheutunut vain vähän ongelmia. Lentokentän toiminnasta vastaava Finavia kertoo, että seuraava jännitysmomentti on varsinaisen mielenilmauksen aikana eli kello yhdestätoista kolmeentoista. Silloin töitä pyritään hoitamaan esimiesvoimin.

---------------------------------------------------------------------------------------------------------------------------------------

Viime viikolla puhuin siitä, miten elämä ei tottele suunnitelmiamme. Se ei kuitenkaan tarkoita sitä, ettenkö uskoisi unelmiin ja niiden toteutumiseen. Päinvastoin. Monet suurista unelmistani ovat toteutuneet ja ne tuottavat minulle suurta iloa nykyäänkin, päivittäin. Viime vuosina toteutuneiden unelmieni listalla ovat muun muassa suuri muutos työelämässä, tupakoinnin lopettaminen, moottoripyöräilyn opetteleminen ja itseni treenaaminen rapakuntoisesta sohvaperunasta keskivertoa parempikuntoiseksi aikuiseksi. Tarkoitus ei ole kehuskella. Unelmien toteutumisessa kun ei ole kyse keijujen ripottelemasta taikapölystä tai poikkeuksellisesta tahdonvoimasta. Minä olen pohjimmiltani mukavuudenhaluinen ihminen. Olen vain sattumalta löytänyt toimivan reseptin, jota olen toistanut unelmasta toiseen. Koska se ei ole salaisuus, kerron sen nyt sinulle. Kaikki alkaa unelmoinnista. On unelmoitava suurista ja suuresti. Sellaisista asioista, jotka alkuun saattavat tuntua jopa saavuttamattomilta. Kun jokin toive palaa ajatuksiini päivästä, viikosta ja kuukaudesta toiseen, ymmärrän, että sitä minun pitää tavoitella. Tämä haudutteluvaihe voi kestää kauan, vuosiakin. Hyvin haaveiltu on puoliksi tehty. Mielellä on valtava voima. Siinä vaiheessa, kun pystyn kuvittelemaan unelmani toteutumisen, tiedän, että se on mahdollista myös oikeasti. Unelman pitää pelottaa, houkuttaa ja jännittää samaan aikaan. Kun kuvittelee sen toteutuvan, vatsan pohjasta vihlaisee. Aidon ja oman unelman tunnistaa siitä, että se herättää itsessä suuria tunteita. Unelma ei tunnu koskaan samantekevältä.

---------------------------------------------------------------------------------------------------------------------------------------

**Part 2**

Millanen kämppä, kerro lisää ihmeessä siitä et..

Joo siis mulla on sellainen kimppa kämppä kaksio, 62 neliöö siellä, se on ihan tosi kiva. Ainoo on vaan et nyt me ollaan itseasiassa muuttamassa mun kämppiksen kanssa, että tota että ens kuun lopussa niin mä muutan siihen ihan toiselle puolelle puistoa. Siellä on sellainen kiva yksiö varattuna mulle. Mut et jään kuitenkin tonne Arabianrantaan asustelee, ku siihen on niin ihastunut. Mut et joo et muutto edessä.

Tuleeks sun kämppis mukaan?

Ei tuu, se muuttaa itseasiassa sen poikaystävän kanssa yhteen.

Joo

Niin mä muutan sit ihan yksin. Mut se on tosi kiva, koska mä oon asunu niin monta vuotta noiden kavereiden kanssa. Niin sit on vähän kiva kokeilla omii siipii ja et miten asuminen luonnistuu yksiössä.

No eiköhän se luonnistu ihan hyvin.

Mm..

Ei siin mitää.

Onks sulla ollu kimppakämppiä vai yksiöitä?

Joo mä asun itseasiassä tälläkin hetkellä, se on sellainen kaverikämppä.

Okei.

Et siinä on tavallaan kaks samanlaista soluhuonetta ja sitten yhteiset keittiöt ja muut, mut et kämppis on oma kaveri, niin se on toiminut tosi hyvin et.

Mm, se on kyl tosi hyvä ratkasu.

Joo niin sit et me opiskellaan molemmat samaa alaakin vielä niin sit voidaan aina pähkii samaan… samaan aikaan niit tehtävii

Mmm.

---------------------------------------------------------------------------------------------------------------------------------------

Kesti kauan aikaa ennen kuin ymmärsin mistä hän tuli. Pikku prinssi, joka itse kyseli minulta paljon, ei tuntunut koskaan kuulevan minun kysymyksiäni. Sattumoisin lausutut sanat paljastivat minulle asian vähitellen. Niinpä hän kysyi nähtyään ensimmäisen kerran lentokoneeni (en piirrä lentokonettani, se olisi minulle aivan liian vaikea tehtävä):

”Mikä kapine tuo on?”

”Ei se ole mikään kapine, se lentää. Se on lentokone. Minun lentokoneeni.”

Olin oikein ylpeä, kun sain kertoa hänelle, että minä lensin.

Siihen hän huudahti: ”Ihanko totta! Olet siis pudonnut taivaasta!”

”Niin”, vastasin vaatimattomasti.

”Ai, sehän on hassua.” Ja pikku prinssi purskahti iloiseen nauruun, joka ärsytti minua tavattomasti. Olin odottanut, että hän suhtautuisi vakavasti onnettomuuteeni. Sitten hän lisäsi:

”Sinäkin tulet siis taivaasta, miltä tähdeltä sinä olet?”

Aloin vähän aavistella, miten hänen läsnäolonsa oli selitettävissä ja kysäisin nopeasti, ”Sinä siis tulet toiselta tähdeltä?” Mutta hän ei vastannut minulle. Katsellen lentokonettani hän pudisti päätään.

”Selvä juttu, ei tuommoisella kapineella voi tulla kovin kaukaa”. Ja hän vaipui syviin ajatuksiin.

Sitten hän veti lampaani taskustaan ja syventyi tarkastelemaan aarrettaan. Voitte kuvitella, miten tämä vihjaus toisiin tähtiin oli kiihottanut uteliaisuuttani. Koitin saada lisää selvitystä asiaan. ”Mistä sinä tulet pikkumies? Missä on kotisi? Minne tahdot viedä lampaani?”

Hiljaisen mietiskelyn jälkeen hän vastasi: ”Oli hyvä, että annoit minulle laatikon. Sitä voi mainiosti pitää lampaan talona yöllä.”

”Niin voikin ja jos olet kiltti, niin annan sinulle myös nuoran ja paalun, jotta voit sitoa lampaan päiväksi”. Ehdotukseni tuntui jotenkin kummastuttavan pikku prinssiä.

”Sitoa kiinni. Mikä hassu ajatus.”

”Mutta ellet pane sitä kiinni, niin se voi mennä vaikka minne ja eksyä.”

Ystäväni purskahti uudelleen nauruun.

”Minne kummaan se voisi mennä?”

”Minne vain. Suoraan eteenpäin…”

Silloin pikku prinssi huomautti vakavasti: ”Se ei merkitse mitään, minun tähdelläni on kaikki niin pientä” ja luulen että hän lisäsi vähän surullisena: ”Suoraan eteenpäin ei voi mennä kovin kauan.”

---------------------------------------------------------------------------------------------------------------------------------------

Hei Paula moikka!

Moi!

Sä olit sillon kesällä siellä Prahassa etkö ollutkin?

Joo olin.

Olikse hyvä reissu?

Siis oli se kyllä, joo ei oo tullukaan puheeks se.

Joo

Siis kyllä. Me oltiin viikko ja kyllä siinä niinku ehti.. ehti mun mielestä oleelliset nähä.

Joo. Mitä sä tykkäsit siit kaupungista yleisesti?

No siis se niin ku se kaupunki sillee just nimeomaan niin ku siellä vaan käveleskeleminen on ehkä just se paras juttu.

Mmm

Se on tosi viehättävä historiallisesti ja sit just ne sillat ja se joki ja... Tai mä luulen et se on niinku se paras juttu Prahassa, ihan vaan se ku kattelee ympärilleen ja...

Joo siis sehän on aika vanha kaupunki eikö ookkiin?

No joo siis on joo. Ja just semmonen niinku Keski-Eurooppalainen historiallinen kaupunki, missä on.. on joki menee läpi ja on hienoja historiallisia siltoja ja sit on joku iso katedraali ja ja..

Joo

Just semmonen.

Joo. Mä olin itseasiassa kans vuos sitten Prahassa…

Aijaa!

Ja siellä jotekin hämmentävää oli se, että kun ne tiet ei sillä tavalla oo suoraan asemakaavaan niinku.

Siis voi visti, joo, siis mä eksyin niin monta kertaa siellä! Kun ne kaikki tiet kulkee niiden liikenneympyröiden suuntaisesti, semmosia isoja kehiä.

Joo, joo. Se oli ihan käsittämätöntä. Mä viel ehin puhua mun kaverille sillä matkalla että mä on niin hirvittävän hyvä lukemaan karttaa..

Voi ei!

Ja mä vein meitä koko ajan täysin väärille kaduille. Ja se tuntu multa niinku siltä että ne kadut vaihtais paikkaa siinä vaiheessa…

Joo

…kun me vähän edetään.

Joo, joo, mulla oli toi ihan sama tunne ja poikaystävä kuittaili et sä et osaa kyllä ollenkaan suunnistaa

Joo joo, mut se on kyl tosi kiva paikka.

---------------------------------------------------------------------------------------------------------------------------------------
